# Evaluation of Machine Learning-Assisted Directed Evolution Across Diverse Combinatorial Landscapes

**DOI:** 10.1101/2024.10.24.619774

**Authors:** Francesca-Zhoufan Li, Jason Yang, Kadina E. Johnston, Emre Gürsoy, Yisong Yue, Frances H. Arnold

**Author notes:** Corresponding Authors: Frances H. Arnold, Yisong Yue. Present address: Merck & Co., Inc., South San Francisco, CA 94080. Present address: Department of Biosystems Science and Engineering, ETH Zürich, Schanzenstrasse 44, 4056 Basel.

## Abstract

Various machine learning-assisted directed evolution (MLDE) strategies have been shown to identify high-fitness protein variants more efficiently than typical wet-lab directed evolution approaches. However, limited understanding of the factors influencing MLDE performance across diverse proteins has hindered optimal strategy selection for wet-lab campaigns. To address this, we systematically analyzed multiple MLDE strategies, including active learning and focused training using six distinct zero-shot predictors, across 16 diverse protein fitness landscapes. By quantifying landscape navigability with six attributes, we found that MLDE offers a greater advantage on landscapes which are more challenging for directed evolution, especially when focused training is combined with active learning. Despite varying levels of advantage across landscapes, focused training with zero-shot predictors leveraging distinct evolutionary, structural, and stability knowledge sources consistently outperforms random sampling for both binding interactions and enzyme activities. Our findings provide practical guidelines for selecting MLDE strategies for protein engineering.

## Introduction

Engineered proteins are indispensable across myriad applications, serving as effective therapeutics to combat diseases, non-toxic agents to enhance crops, and green biocatalysts to synthesize chemicals.^1^ The development of such useful proteins often involves directed evolution (DE), a method for accumulating beneficial mutations using iterations of mutagenesis and functional assessment by selection or screening.^2–4^ DE is an empirical, greedy hill climbing process on a high-dimensional fitness landscape that maps protein sequence to function (Figure 1a).^5,6^ Despite its widespread use, DE remains time-consuming and resource-intensive: screening is expensive, and multiple rounds of mutation and screening may be needed to generate the desired improvements.

**Figure 1.**
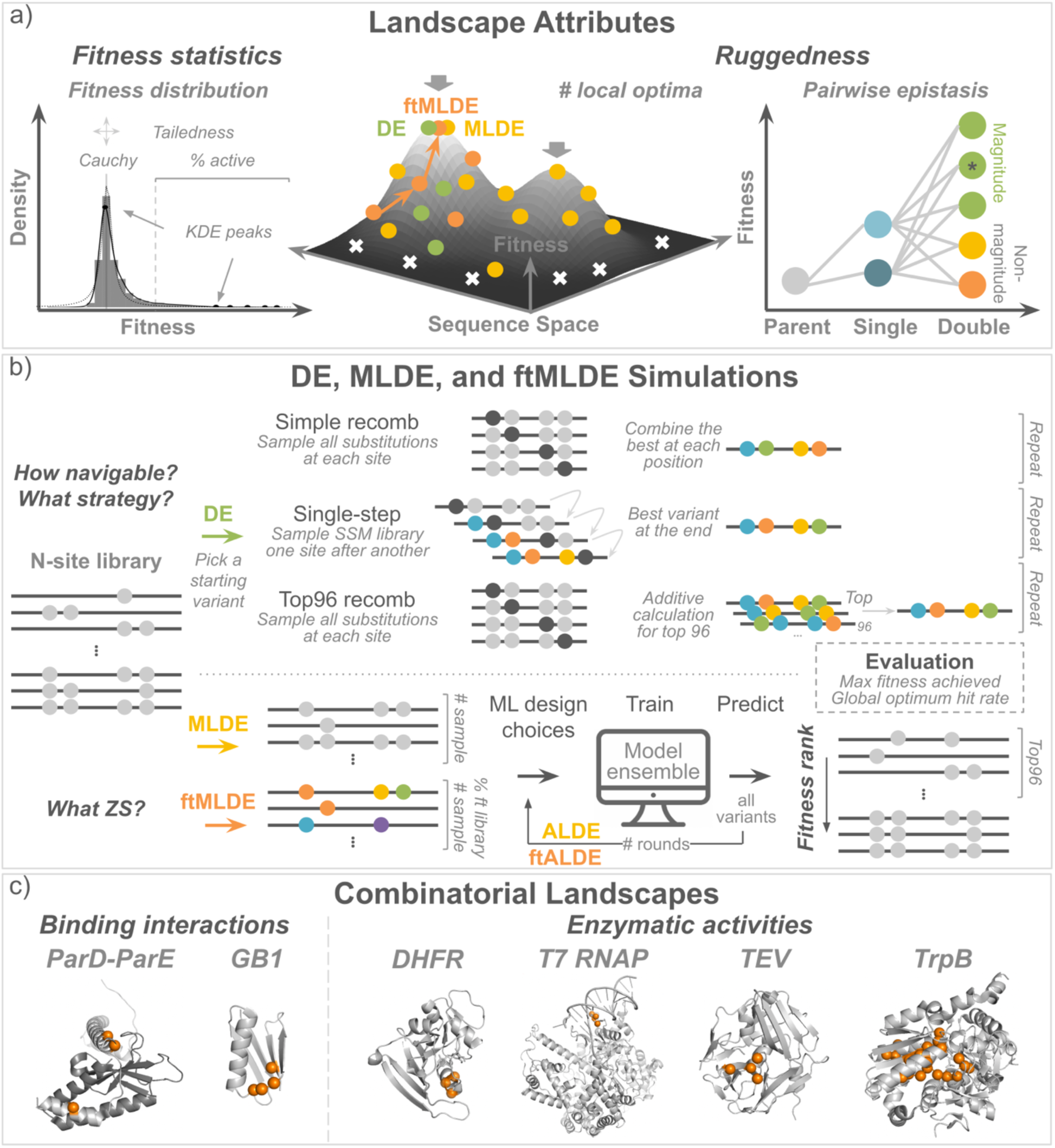
Summary of landscape attributes, simulations, and combinatorial landscapes. a) Landscape attributes include fitness statistics (percent of active variants, tailedness, location of Cauchy peak, and number of peaks for kernel density estimation) and ruggedness (number of local optima and percent of non-magnitude pairwise epistasis). b) Various in silico simulations include three types of DE (simple recomb, single-step, and top96 recomb), MLDE, multiple rounds (ALDE), and focused training for both MLDE (ftMLDE) and ALDE (ftALDE) (Methods). c) Combinatorial landscapes studied, categorized by the number of targeted sites (three and four) and function types (binding interactions and enzyme activities) on six protein systems (ParD-ParE toxin-antitoxin, GB1 immunoglobulin binding, dihydrofolate reductase, T7 RNA polymerase, TEV protease, and tryptophan synthase).^37,40,42–44^ To keep the main text relevant to the majority of campaigns, we focused on libraries with at least 1% active variants, while the remaining landscapes are detailed in the supplemental information.

Fitness landscapes are more rugged and difficult to traverse when rich in epistatic, or non-additive, effects of amino acid substitutions.^7,8^ Epistasis is often observed between mutations in close structural proximity^9^ and is enriched at binding surfaces or enzyme active sites, due to direct interactions between residues, substrates, and/or cofactors.^8^ Protein engineers frequently target mutations to these interacting sites to enhance a function, often using simultaneous site-saturation mutagenesis (SSM) to make libraries in which the targeted amino acids are mutated to many or all 19 other possible amino acids.^10^ Combining the beneficial mutations found at these sites often reveals epistatic effects. For example, beneficial mutations in the context of the initial sequence may not be beneficial in combination with other mutations. Therefore, epistasis can present a significant challenge for DE.^3^

Compared to DE, machine learning-assisted DE (MLDE) has shown promise for exploring a broader scope of sequence space and more effectively navigating epistatic landscapes.^11–13^ MLDE utilizes ML models trained on sequence-fitness data to capture non-additive effects. The trained models can then be used to predict high-fitness variants across the entire landscape in a single round^11,12,14^ or iteratively in an active learning (ALDE) fashion.^15–17^

The choice of the training set can greatly influence the performance of the ML models. One can randomly sample the full combinatorial space for training the model (MLDE) or alternatively do focused training (ftMLDE)^12^ by selectively sampling to avoid low-fitness variants. In the latter, the quality of the training set can be biased toward more-informative variants with zero-shot (ZS) predictors to reach high-fitness variants more effectively. ZS predictors estimate protein fitness without the need for experimental data: they are instead based on prior assumptions and leverage auxiliary information, such as protein stability calculations, evolutionary data, or structural information.^18–25^

Although ML in protein engineering has been demonstrated in different case studies,^11,26–36^ most MLDE^11,15,16^ and ftMLDE^12^ studies on epistatic landscapes have been benchmarked against a single dataset on the B1 domain of an immunoglobulin-binding protein G (GB1).^37^ Thus, two key issues persist: first, the effectiveness of different MLDE strategies on proteins with complex functions, such as enzymes, remains uncertain and, second, the principles that guide successful use of MLDE strategies across diverse protein properties are not understood. Furthermore, despite a growing array of ZS options,^25^ there is no definitive guideline for selecting predictors for a given application. This is particularly true for combinatorial epistatic landscapes.^23,38^

Recent experimental studies^39–44^ provide a wealth of data on a broader array of protein fitness landscapes, enabling us to start establishing best practices and generalizable guidelines for practitioners working with various proteins. To contextualize the benefits of MLDE, ALDE, and focused training, we conducted a comprehensive study of 16 diverse combinatorial protein fitness landscapes. They span six protein systems and two function types (binding and enzyme activity). Consisting of variants that are simultaneously mutated at three or four residues, these landscapes vary in landscape attributes, such as statistical measures (including the number of active variants and fitness distribution properties) as well as ruggedness (a measure of the prevalence of fitness interactions among variants,^45^ including pairwise epistasis and the number of local optima). Specifically, this study focuses on two questions: (1) When do MLDE, multiple rounds (such as in ALDE), and focused training offer a significant advantage compared to DE? (2) How can we best select and utilize the ZS predictor(s) for focused training?

## Results

### Overview of landscapes

For this study, we selected 16 experimental combinatorial landscapes covering a range of binding interactions and enzyme activities (Table 1 and S1).^37,40–44^ All landscapes feature mutations at binding interaction points, in active sites, or at positions previously shown to modulate fitness, all of which are often targeted for engineering tasks (Figure 1c). For binding, we examined two three-site bacterial toxin-antitoxin ParD-ParE landscapes^40^ and the GB1 landscape for immunoglobulin binding.^37^ For enzyme activity, we analyzed a three-site dihydrofolate reductase (DHFR) landscape,^41^ a three-site T7 RNA polymerase landscape,^42,43^ a four-site TEV protease landscape,^42,43^ and ten three-or four-site landscapes of the thermostable β-subunit of tryptophan synthase (TrpB).^44^

**Table 1.**
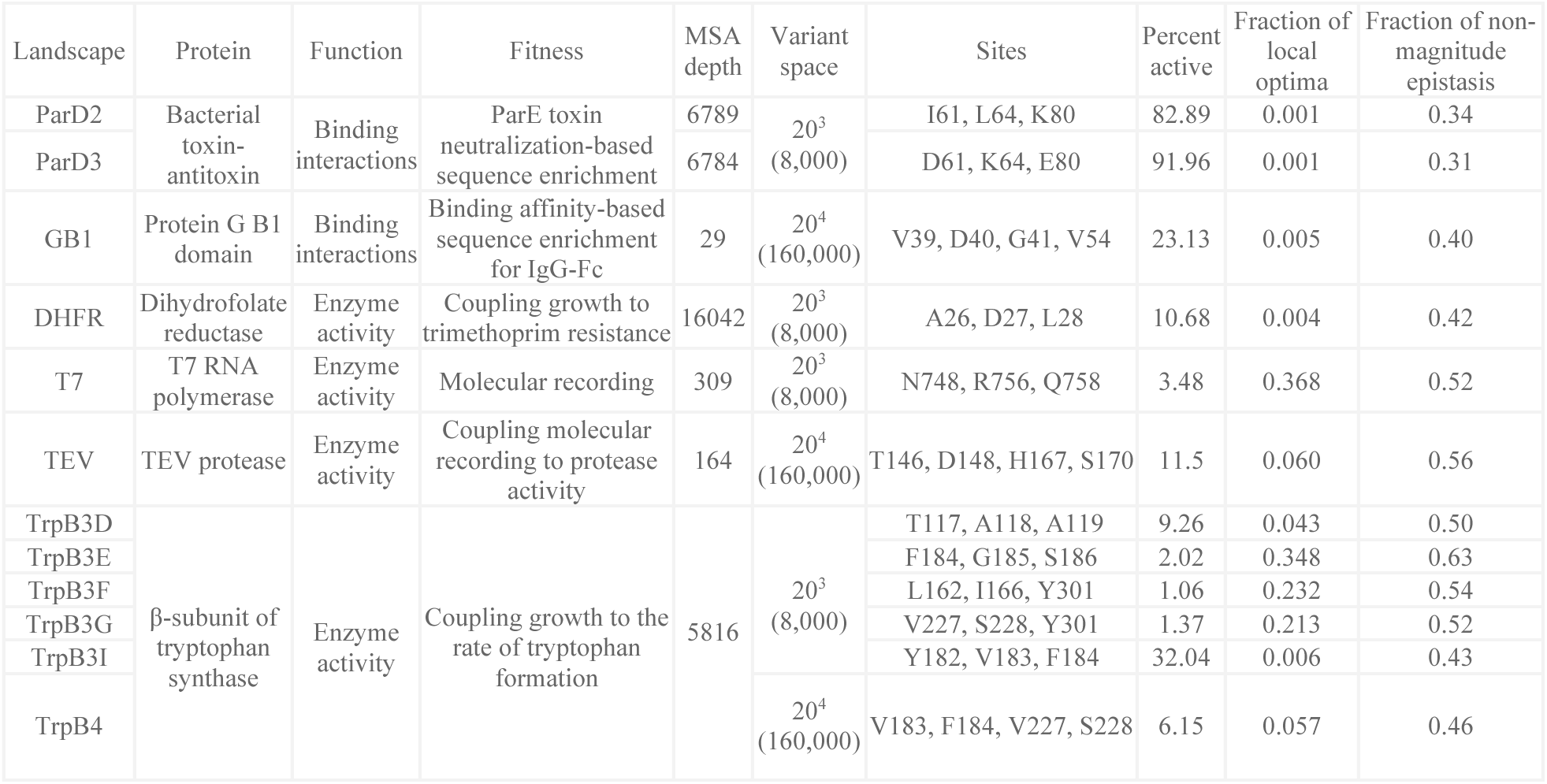
Combinatorial landscapes analyzed in the main text (for full details, see Table S1). ^37,40–44^.

Due to the frequent misalignment between theoretical landscape modeling and experimental applications, we selected two groups of empirical and interpretable attributes to characterize the landscapes: fitness statistics (which do not incorporate sequence information) and landscape ruggedness (which involves mapping sequences to fitness) (Figure 1a; Methods).^46–53^ We used the following statistics to indirectly infer the complexity of the fitness landscape: the percentage of active variants, the fitness value corresponding to the Cauchy peak location, the kurtosis of the fitness distribution (“tailedness”), and the number of kernel density estimation (KDE) peaks (Methods). The Cauchy distribution is known for its heavy tails. We sought to use the fitness corresponding to its peak location as a landscape attribute to capture the majority of the variant finesses. KDE is a non-parametric method for estimating the probability density function of fitness distribution. KDE does not assume any specific underlying distribution for fitness and is useful for smoothing out noise. We reasoned that the number of KDE peaks, reflecting the distribution modalities of fitness, could serve as a proxy for the underlying landscape navigability, which impacts the outcome of DE.

To quantify ruggedness, which poses navigation challenges for DE,^47,49^ we included the number of local optima and the percentage of pairwise non-magnitude epistasis (Figure 1a). We defined a local optimum as a variant that possesses higher fitness than all its active neighbors differing by a single amino acid substitution (Methods).^41,44,46,47,49^ Recent studies have also highlighted the impact of various types of epistasis on DE^3,44^ and emphasized that the majority of epistasis is pairwise.^54^ Thus, we also included the amount of non-magnitude pairwise epistasis (conditional or impossible for DE to navigate) as a relevant landscape attribute (Methods).

### All MLDE strategies consistently outperform DE, particularly as landscape navigability decreases

We next assessed how landscape attributes influence the efficacy of protein engineering strategies. Specifically, we evaluated the outcomes of a protein engineering campaign using two metrics: (1) “average maximum fitness achieved,” which is the fitness of the final variant achieved by each method on average and (2) “fraction reaching the global optimum”, which measures how frequently the true maximum fitness is reached. We explored these measures across MLDE, ALDE, focused training, and three different DE strategies. The DE strategies are summarized as “recomb”, a recombination of the best SSM variant at each site (19 × *n_site_* + 2 samples, including the initial and final variant); “single-step”, an iterative process starting from any site with subsequent variants built on the best variant found (19 × *n_site_* + 1 samples, including the initial variant); and “top96 recomb”, where SSM is performed at each position, all substitution combinations are calculated based on additive recombination, and the top 96 variants are selected (19 × *n_site_* + 97 samples, including the initial and 96 variants, where 96 is the number of wells in plates commonly used for screening; Figure 1b; Methods).^11,12,44^ MLDE and ftMLDE trained an ensemble of models, employing either random or ZS predictor-guided training sample selection. The trained models were then used to predict fitness for all variants, where the top 96 predicted variants were then used for evaluation (Methods).^12^ ALDE divided the total sample size into multiple rounds, with each subsequent round of sampling guided by the uncertainty quantified in its previous round (Figure 1b; Methods).^17^ Similar to how ftMLDE improved upon MLDE, ftALDE used ZS predictors to select a more informative initial training set, instead of the random sampling used in ALDE.

Considering the variability in throughput and expense of experimental screens, we explored a range of total number of variants screened (total sample size), from 120 to 2,016 samples (Figure 2a; Table S2). On average across landscapes, MLDE (dashed light blue line) required merely 48 training samples to outperform “recomb” DE and 96 to surpass “single-step” DE for both metrics. It took 96 training samples for MLDE to match the average maximum fitness and 384 to achieve a comparable fraction reaching the global optimum as the most competitive DE strategy, “top96 recomb”. By incorporating various ZS predictors, ftMLDE (solid dark blue line) consistently outperformed MLDE with random sampling (showing a 4–12% improvement in average maximum fitness achieved for up to 960 training samples and a 9–77% improvement in fraction reaching the global optimum across all training sample sizes, Table S3); ftMLDE achieved the same levels of average maximum fitness and global optimum fraction as MLDE but with fewer training samples required (Figure 2a). These results suggest that MLDE can identify high-fitness variants more effectively than DE, and focused training with ZS predictors can further improve performance compared to MLDE with random sampling.

**Figure 2.**
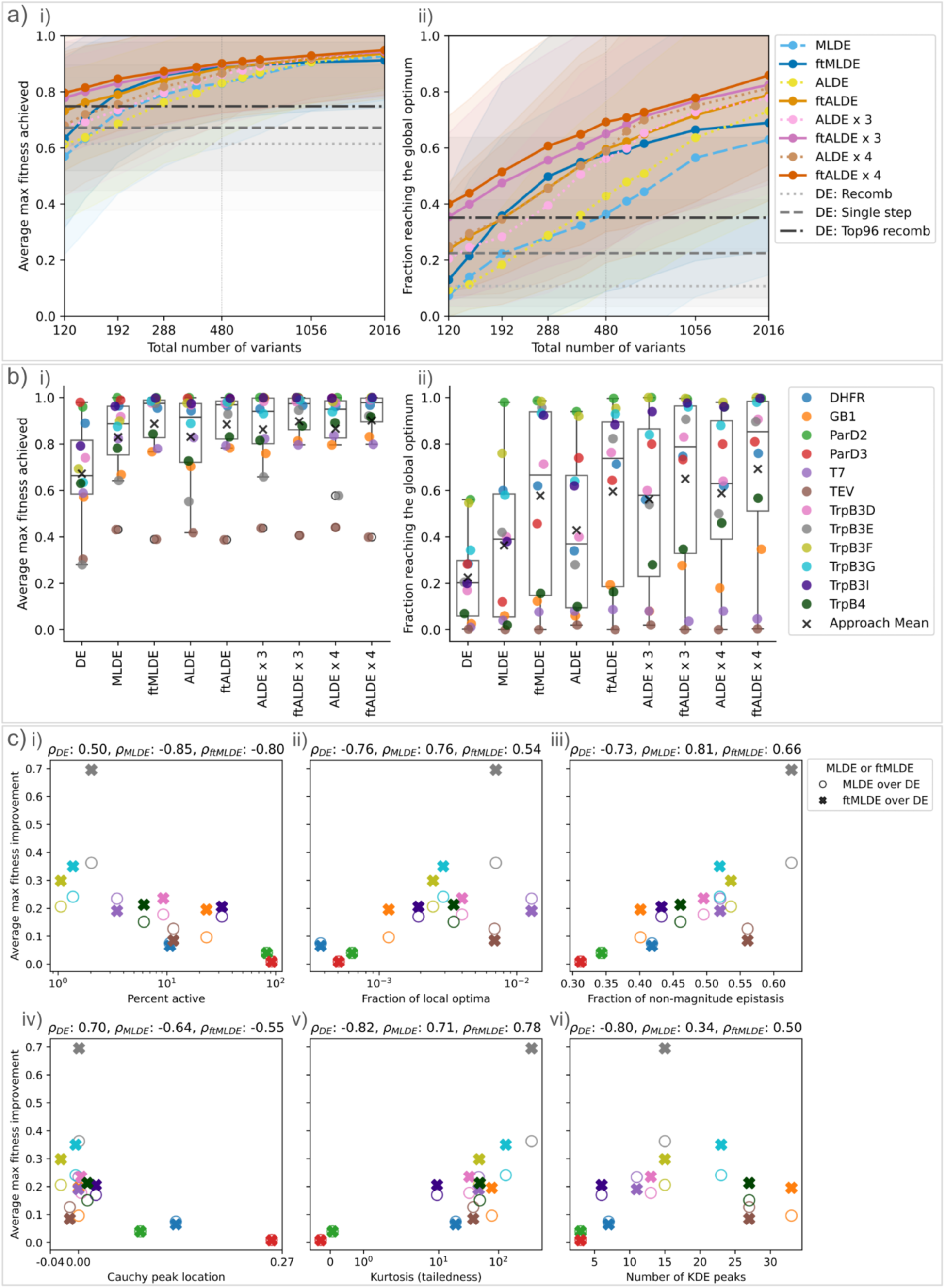
Performance and correlations of MLDE, ALDE, and focused training compared with DE and six landscape attributes. a) Comparison of DE, MLDE, ALDE, ftMLDE, and ftALDE performance, averaged across 12 landscapes with at least 1% active variants. Shading indicates standard deviation. Performance is shown for i) the maximum fitness achieved and ii) the fraction of campaigns reaching the global optimum, for different numbers of training samples. Three DE methods were included: simple recomb, single-step, and top96 recomb. The vertical line marks a total sample size of 480 (e.g. 384 sampled variants for training and top 96 predicted variants for testing, or four rounds of ALDE each with 120 variants) that the following results expand on. See Figure S1 for landscapes with fewer than 1% active variants. b) Single-step DE, MLDE, ALDE, and focused training results broken down by landscape, with a total sample size of 480 for both metrics. See Figure S2 for landscapes with fewer than 1% active variants. c) Spearman’s ranking correlation of ML strategy performance improvement (the average maximum fitness of the top 96 predicted variants by MLDE and focused training over single-step DE, y-axis) with six landscape attributes (x-axis): i) percentage of active variants, ii) fraction of local optima (normalized to the number of variants measured), iii) fraction of non-magnitude epistasis, iv) Cauchy peak location, v) kurtosis (tailedness), and vi) number of kernel density estimation (KDE) peaks. See Figure S3 for different rounds of ALDE and ftALDE. For each MLDE and ftMLDE experiment, boosting models were trained on 384 random samples from the entire or ZS-focused library using one-hot encoding, with five-fold cross-validation. For each ALDE and ftALDE experiment, boosting ensembles with greedy acquisition function were trained with 240, 160, or 120 samples per round for two, three, or four rounds in total, respectively. The top 96 predicted variants were evaluated. Each ML experiment was averaged across 50 replicates (Methods). ftMLDE and ftALDE performance were further averaged across six ZS predictors (details in the next section). DE simulations started from all active variants (Methods).

Next, we compared ALDE (MLDE with multiple rounds of training and testing guided by uncertainty quantification) to MLDE (a single round of training and testing, equivalent to two rounds) with the same total number of variants screened. With two rounds, ALDE (dotted bright yellow line) began to outperform MLDE (dashed light blue line) after 480 total samples for average maximum fitness achieved and 288 total samples for fraction reaching the global optimum but did not outperform ftMLDE (solid dark blue line) until 1,056 samples for both metrics. With four rounds, ALDE (dotted light brown line) matched or exceeded ftMLDE performance. With focused training, ftALDE (solid orange line) matched or surpassed ftMLDE with the same number of rounds and showed further improvement with additional rounds (Figure 2a). However, for libraries with fewer than 1% active variants, even four rounds of ALDE (without focused training) consistently underperformed compared to ftMLDE (Figure S1). Our observations underscore the utility of focused training using ZS predictors, enabling MLDE to match multi-round ALDE performance and offering further improvement to ALDE.

Given the large standard deviations in the performance of different approaches across landscapes, we examined how each approach performed on individual landscapes and found that some landscapes exhibited more significant improvements than others (Figure 2b and S2). We first quantified the improvements of ML strategies over single-step DE and found that ML strategies offered a greater advantage on landscapes which were more challenging for DE to navigate. To better understand when DE struggled, we then calculated six different attributes to provide insights into landscape navigability (Figure 2c; Methods). Specifically, the mean maximum fitness achieved by DE correlated positively with the fraction of active variants (Figure 2c–i) and the fitness distribution’s Cauchy peak location (Figure 2c–iv), indicating improving navigability by DE. Consequently, the improvements resulted from all ML methods were anti-correlated with percent active (Figure 2c–i) and Cauchy peak location (Figure 2c–iv). The amount of kurtosis (tailedness) and number of KDE peaks of the fitness distribution hindered DE navigability (Figure 2c–v and vi). ML methods thus improved performance most significantly for landscapes with high tailedness and more KDE peaks. Similarly, increased landscape ruggedness decreased DE navigability, yielding greater benefit of using ML methods over DE for such landscapes. DE navigability was anti-correlated with the fraction of local optima (Figure 2c–ii) and the fraction of non-magnitude epistasis (Figure 2c–iii), and thus the net improvement of ML methods over DE was positively correlated with both higher fractions of local optima (Figure 2c–ii) and higher fractions of non-magnitude epistasis (Figure 2c– iii). Indeed, ftMLDE demonstrated the most substantial performance improvements (3.5-fold) for one of the least navigable landscapes (TrpB3E; Table S4). The performance improvements from different rounds of ALDE and ftALDE were also correlated with landscape navigability defined by the six attributes (Figure S3; Table S4).

### ZS predictors provide orthogonal priors on protein fitness that improve focused training performance

Next, we sought to understand the effectiveness of different ZS predictors for fitness prediction and their ability to improve ftMLDE and ftALDE performance across landscapes. ZS predictors could be useful for (1) effectively ranking variants to sample the fittest mutants (measured by Spearman’s correlation, Methods) and (2) filtering out non-viable variants, especially in landscapes dominated by inactive variants (measured by ROC-AUC, Methods). To evaluate the effectiveness and limitations of various ZS predictors under different assumptions and priors, we selected six distinct predictors across two axes: calculation vs. learning-based and sequence vs. structure-based. These predictors include Hamming distances, EVmutation, ESM, ESM-IF, CoVES, and Triad (Figure 3a; Methods).^18–22,55^ To differentiate our work from comprehensive ZS predictor benchmarks that are largely limited to measuring the effects of single amino acid substitutions,^25^ we emphasized their utilities for focused training applications in epistatic landscapes.

**Figure 3.**
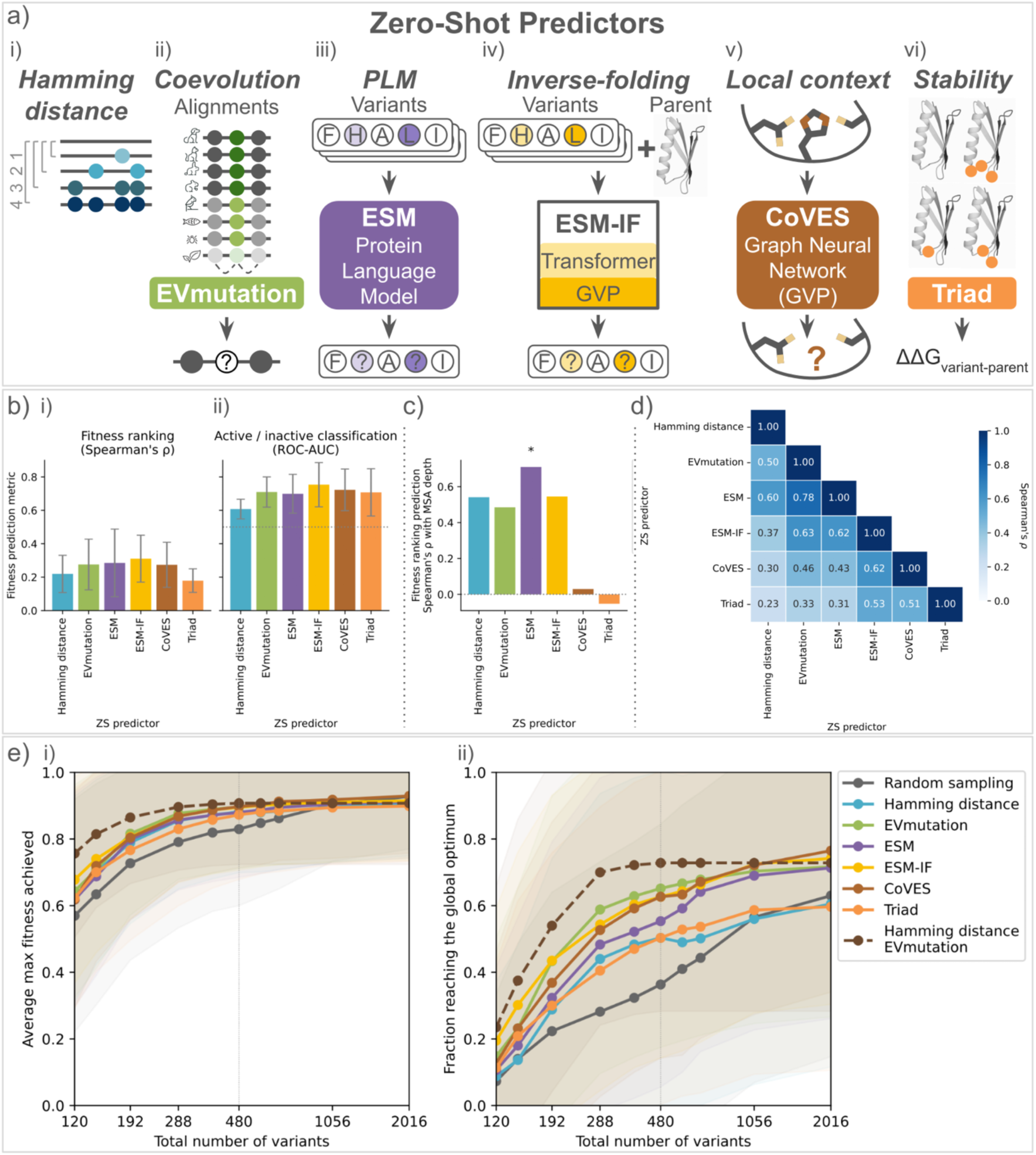
Summary of different ZS predictors and their impacts on focused training across landscapes. a) Six ZS predictors: i) Hamming distance, ii) EVmutation (coevolutionary conservation),^18,56^ iii) ESM (mutant likelihood from pretrained protein-language model),^19^ iv) ESM-IF (mutant likelihood from pretrained inverse folding models based on sequence and structure information),^20^ and v) Triad (mutant stability ΔΔG).^12,55^ b) ZS predictor performances in terms of variant fitness ranking correlation (Spearman’s ρ) and active/inactive classification (ROC-AUC) across 12 landscapes with at least 1% active variants. Error bars indicate standard deviation. Dotted gray line indicates random classification. c) Fitness values ranking Spearman’s correlation with MSA depth for each ZS predictor. Statistical significance (p-value <0.05) is indicated as * (Table S5). Dotted gray line indicates no correlation. d) Pairwise Spearman’s correlation of six ZS predictors averaged across 12 landscapes. e) Performance of focused training with different ZS predictors including the best Hamming distance ensemble, averaged across 12 landscapes. Shading indicates standard deviation. It assesses i) the maximum fitness achieved and ii) the fraction reaching the global optimum in relation to the number of samples used by ftMLDE. For each MLDE and ftMLDE experiment, boosting models were trained on 384 random samples from the entire or ZS-focused library using one-hot encoding, with five-fold cross-validation. The top 96 predicted variants were evaluated. Each ML experiment was averaged across 50 replicates (Methods). The vertical line marks a total sample size of 480 (e.g., 384 sampled variants for training and top 96 predicted variants for testing). ALDE and ftALDE results see Figure S7. For landscapes with fewer than 1% active variant, see Figure S8, S9, and S12.

As a baseline, we used the Hamming distance as a ZS predictor, which counts the number of amino acid substitutions from the parent, a variant already exhibiting some activity. By setting a Hamming distance threshold, we essentially confined the sampling space to the vicinity of the parent to enrich the training set with more viable variants on average, as most mutations are deleterious.^5^ Indeed, we observed that the Hamming distance (indicated in blue) showed a weak correlation with fitness ranking (Figure 3b–i) and classified active/inactive variants better than random (Figure 3b–ii) as a ZS predictor. Notably, Hamming distance relied on the parent defined by the authors of each landscape, rather than a randomly sampled active variant as in the DE simulations. Since these landscapes were designed to have variant activity levels both higher and lower than the parent, the parent and its neighboring variants were likely to be more fit and active than those around a randomly selected active variant (Figure S4). Although the landscape parent was one of the most active, leading to strong Hamming distance performance, Hamming distance still outperformed random predictions in fitness ranking (Figure S5) and active/inactive classification (Figure S6) on average, when using different active variants as the parent. In the focused training setting, Hamming distance-guided training set sampling outperformed random selection, improving both the average maximum fitness achieved across all total sample sizes (Figure 3e–i) and fraction reaching the global optimum, up to total sample sizes of 1,056 for ftMLDE (Figure 3e–ii) and 480 for ftALDE (Figure S7).

Various ZS predictors can incorporate implicit evolutionary conservation based on the distribution of naturally occurring sequences. The EVmutation score predicts the fitness effect of a given set of substitutions based on conservation and evolutionary couplings through multiple-sequence alignment (MSA).^18,56^ We observed EVmutation (indicated in green) outperformed the Hamming distance for both ranking fitness values (Figure 3b–i) and classifying active/inactive variants (Figure 3b–ii). Moreover, it was one of the best ZS predictors for focused training across all sample sizes on both metrics (Figure 3e). Similarly, protein language models (PLMs) can capture these evolutionary conservations by learning to predict the original identity of masked or corrupted amino acids.^57–63^ The likelihood of filling such amino acids can be thought of as a predictor for different amino acid substitutions given the sequence context.^19^ The ESM score (Evolutionary Scale Modeling, indicated in purple) from one of such state-of-the-art PLMs performed similarly to EVmutation as a ZS predictor for both fitness ranking (Figure 3b–i) and active/inactive classification (Figure 3b–ii). It also did not further improve upon EVmutation in the focused training setting (Figure 3e).

Incorporating structural context can also be useful for ZS predictions. ESM-IF (ESM inverse-folding, indicated in yellow) is an inverse-folding model trained to predict a protein sequence from its backbone atom coordinates, where effects of substituting amino acids can be approximated with the likelihoods of each possible sequence for this reconstruction task.^20^ We observed that ESM-IF was the best ZS predictor for fitness ranking (Figure 3b–i) and active/inactive classification (Figure 3b-ii), but with only slight improvements over other ZS predictors. In the focused training setting, ESM-IF score did offer a consistent advantage over the sequence-only ESM, but only offered a slight advantage over EVmutation at either low or high number of samples (i.e., 120, 144, 1,056, and 2,016, Figure 3e). CoVES (Combinatorial Variant Effects from Structure, indicated in brown) learns to predict a masked amino acid identity from its surrounding atomic-level structural microenvironments but does not account for epistasis.^21^ Compared to ESM-IF, we observed that CoVES was a slightly less effective ZS predictor for fitness estimation (Figure 3b) and in the focused training setting (Figure 3e), but it was still one of the most effective predictors for improving over random sampling.

An alternative local structure-based ZS score utilizes physics-informed stability calculations. Stability is an important prior for protein function, as an unfolded or misfolded protein will be less likely to be functional.^12,22^ The Triad score estimates mutant stability by calculating the change in its free energy of folding relative to the parent (ΔΔG) using a Rosetta energy function.^12,55^ While Triad (indicated in orange) was the weakest predictor for variant fitness ranking (Figure 3b–i), it classified active/inactive variants fairly well (Figure 3b–ii) as a ZS predictor. Triad-guided training set sampling outperformed random selection in the ftMLDE setting, up to a total sample size of 1,056 for both metrics (Figure 3e). In the ftALDE setting, it outperformed random selection up to a total sample size of 576 for average maximum fitness and 384 for the fraction reaching the global optimum (Figure S7). The relative differences between ZS predictors in focused training remained consistent across different rounds of ftALDE. However, in libraries with fewer than 1% active variants, these differences were minimized, and all ZS-guided focused training approaches showed a significant advantage over random sampling (Figure S8 and S9).

To facilitate ZS predictor selection and ensembling, we first examined how the fitness ranking performance of ZS predictors correlated with the depth of multiple sequence alignments (MSAs) (Figure 3c; Table S5; Methods). We found that the performance of the physics-based Triad and the structure-only CoVES did not correlate with MSA depth, confirming their independence from evolutionary data. In contrast, the three sequence-based predictors, Hamming distance, EVmutation, and ESM did show correlation with MSA depth. Despite being a hybrid sequence-structure model, ESM-IF captured evolutionary information to a similar extent as EVmutation, likely because over 99% of its structures were predicted from similar sequence databases (the UniRef family).^20^

We then investigated the relationship between different ZS predictors using pairwise correlations (Figure 3d; Methods).^64,65^ Within each modality, sequence-based (Hamming distance, EVmutation, and ESM) or structure-based (CoVES and Triad), all ZS predictors exhibited at least a 0.5 Spearman’s correlation with each other. ESM and EVmutation showed the strongest correlation (Spearman’s ρ = 0.78), suggesting PLMs like ESM may capture similar coevolutionary information as MSAs used by EVmutation.^58,66,67^ ESM-IF showed similar correlations with both the structure-only CoVES (Spearman’s ρ = 0.62) and the sequence-only ESM (Spearman’s ρ = 0.62) and EVmutation (Spearman’s ρ = 0.63). Despite distinct approaches, all four learning-based predictors captured related information. However, Triad had only moderate correlations with the other two structure-based predictors (Spearman’s ρ = 0.5) and weak correlations with the three sequence-based predictors (Spearman’s ρ < 0.4). Hamming distance showed moderate correlations with ESM and EVmutation (Spearman’s ρ = 0.6 and 0.5, respectively) but it was weakly correlated with the structurally inclined predictors (Spearman’s ρ < 0.4). This underscores the orthogonality between learning-based models, a naive protein engineering prior, and a physics-based approach. Thus, we hypothesized that ensembling orthogonal ZS predictors may further enhance focused training by synergizing complementary information sources.

We evaluated if ensembling Hamming distance with other ZS predictors enhanced focused training performance compared to each predictor alone. Prefiltering with a Hamming distance (by restricting variants to those within two amino acid substitutions of the parent sequence) boosted focused training performance from each ZS predictor up to 192 total samples (left vertical gray line in Figure S10). This benefit extended to 480 total samples for ESM and ESM-IF (right vertical gray line in Figure S10) and continued to 1,056 for EVmutation for both average max fitness achieved (Figure 3e–i) and fraction reaching the global optimum (Figure 3e–ii). However, the benefits diminished beyond a total sample size of 480, due to this pre-constrained sampling space. In contrast, ZS ensembles with Triad did not show significant improvement (Figure S11 and S12). This suggests that, despite its orthogonal information, the physics-based Triad predictor offered no further benefit to focused training performance. Similarly, naively ensembling the two top-performing predictors, ESM-IF and EVmutation, yielded no additional improvements (Figure S11 and S12). Our results highlight the advantage of combining Hamming distance with other informative ZS predictors to further enhance focused training performance.

### Landscape and functional attributes affect ZS predictability

We next examined how ZS predictability differed across landscapes, specifically comparing those measuring binding interactions vs. enzyme activities (Figure 4a). All six ZS predictors ranked fitness values substantially better for binding interactions than for enzyme activity (Figure 4a–i), with Triad showing a statistically significant difference (p-value = 0.001, Table S6). The structure-based predictors (CoVES and Triad) were better at classifying active/inactive variants for binding datasets than for enzymatic ones, while the sequence-based predictors (Hamming distance, EVmutation, and ESM) performed better for enzyme activity datasets. Hamming distance showed a statistically significant difference in the context of classification (p-value = 0.042, Figure 4a–ii; Table S6). However, the differences between binding interactions and enzyme activities were no longer statistically significant for any ZS predictors in the focused training setting across different ML strategies (Figure 4b; Table S7 and S8).

**Figure 4.**
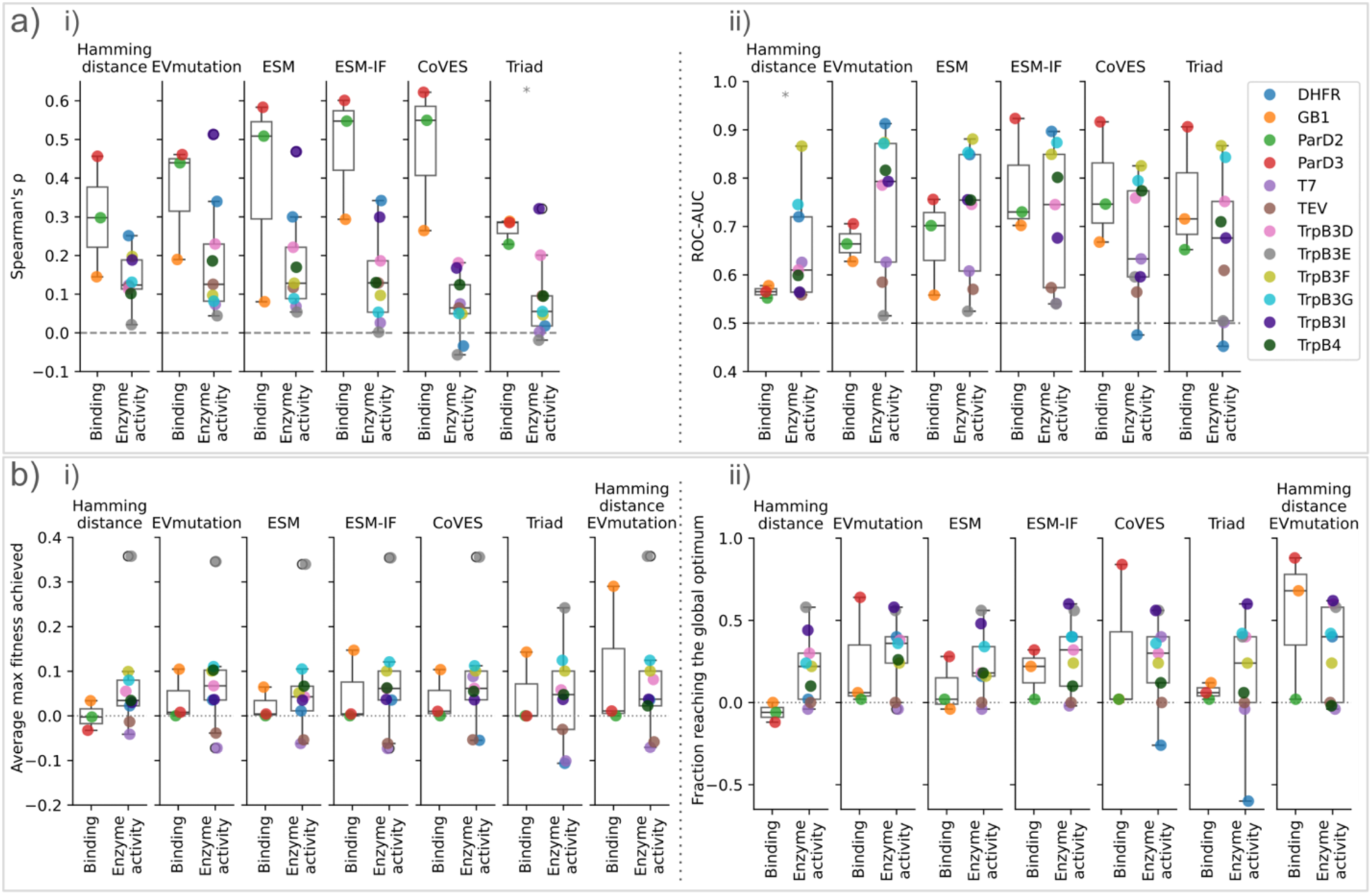
Summary of different ZS scores and their impact on individual landscapes by types of function. a) Six ZS predictor performance for each individual landscape in terms of i) Spearman’s correlation of fitness values ranking and ii) ROC-AUC of active/inactive classification. Random predictions are indicated in horizontal gray dashed lines. Statistical significance (p-value <0.05) is indicated as *. b) A breakdown of the ftMLDE results with a total sample size of 480 from Figure 3e, categorized by six ZS predictors and two functions (binding interactions and enzyme activities) for each landscape. Focused training improvement over randomly sampled training set (MLDE) is quantified by i) average maximum fitness and ii) fraction reaching the global optimum. See supplemental information for landscapes with fewer than 1% active variants (Figure S14 and S15), ftALDE with different rounds (Figure S15 and S16) and ZS predictor impacts on focused training with 192 total samples (Figure S14, S17, and S18).

For 10 out of the 12 landscapes, all six ZS predictors successfully focused the training set to be more informative than random sampling, leading to improved ftMLDE performance for both metrics (Figure 4b). Harder-to-navigate landscapes and the libraries with fewer than 1% active variants benefited more from focused training, provided the ZS predictor for active/inactive variant classification was better than random (ROC-AUC > 0.5, Figure 4, S13, and S14). TrpB3E (indicated in gray), one of the hardest-to-navigate landscapes (Figure 2b) with one of the lowest but still above-random active/inactive variant classification ROC-AUC (Figure 4a–ii), gained the most from all ZS predictors compared to randomly sampled MLDE training sets for both performance metrics (Figure 4b). Similar improvements in hard-to-navigate landscapes were consistently observed when comparing ftALDE with ALDE in the same round, and when considering different total sample sizes for each of the focused training approaches (Figure S15-S18).

A falsely biased training set could negatively impact focused training performance. For DHFR (dihydrofolate reductase, indicated in blue), the structure-based predictions, Triad ΔΔG and CoVES, performed poorly, with worse-than-random active/inactive classification (Figure 4a–ii, dashed gray line) and harmed ftMLDE performance for both metrics (Figure 4b).

## Discussion

Our findings confirmed that all MLDE strategies exceeded or at least matched DE performance across 16 landscapes, with the advantages becoming more pronounced as landscape attributes posed greater obstacles for DE (e.g., fewer active variants and more local optima). ZS predictors, which leverage various prior knowledge, enriched training sets to enable ftMLDE to match multi-round ALDE performance and offered further improvement to ALDE. Overall, our study suggests that MLDE strategies are highly generalizable and can significantly reduce the experimental load of DE, and we present key considerations for the effective deployment of these approaches. We expect that these findings will encourage and facilitate the adoption of ML-assisted directed evolution for efficient protein engineering.

As a general recommendation for the implementation of ML strategies to guide protein engineering objectives, we introduce a decision-making process for selecting a campaign strategy (Figure 5). The first step is to assess whether the landscape is hard-to-navigate. This typically involves a low percentage of active variants and high percentage of pairwise non-magnitude epistasis, which can be inferred from prior knowledge (e.g., the percentage of active variants from single-site SSM experiments) or from the structural proximity of residues of interest to functional (binding or active) sites and to each other. We observed a weak negative correlation (Spearman’s ρ = -0.34) between the percentage of pairwise non-magnitude epistasis (where higher values indicate harder-to-navigate landscapes) and the average pairwise C-alpha distance of mutated residues (the smaller the distance, the closer the central carbon atoms of the two amino acids at the targeted sites, Methods). Next, determine whether there is a “good-enough” ZS prior (optionally ensembling orthogonal predictors), meaning a ZS predictor that can classify active/inactive variants better than random. For hard-to-navigate landscapes without prior information, using a Hamming distance threshold of two (i.e., constructing double-site libraries) can effectively enrich informed variant sampling for the training sets. Additionally, ZS predictor classification performance on single-site libraries can identify predictors that may fail on larger combinatorial libraries as well (Figure S19 and S20; Table S9). For example, CoVES and Triad active/inactive variant classification for DHFR were worse than random for single substitutions (Figure S19) and they both classified the full DHFR landscape worse than random (Figure 4a–i), which ultimately ablated the benefit of focused training (Figure 4a–ii). Finally, consider whether the search space is large (e.g., four-site libraries) and whether the screening budget allows for multiple rounds and/or an increased number of samples per round. A decision tree is provided to assist users in selecting the appropriate strategy (Figure 5).

**Figure 5.**
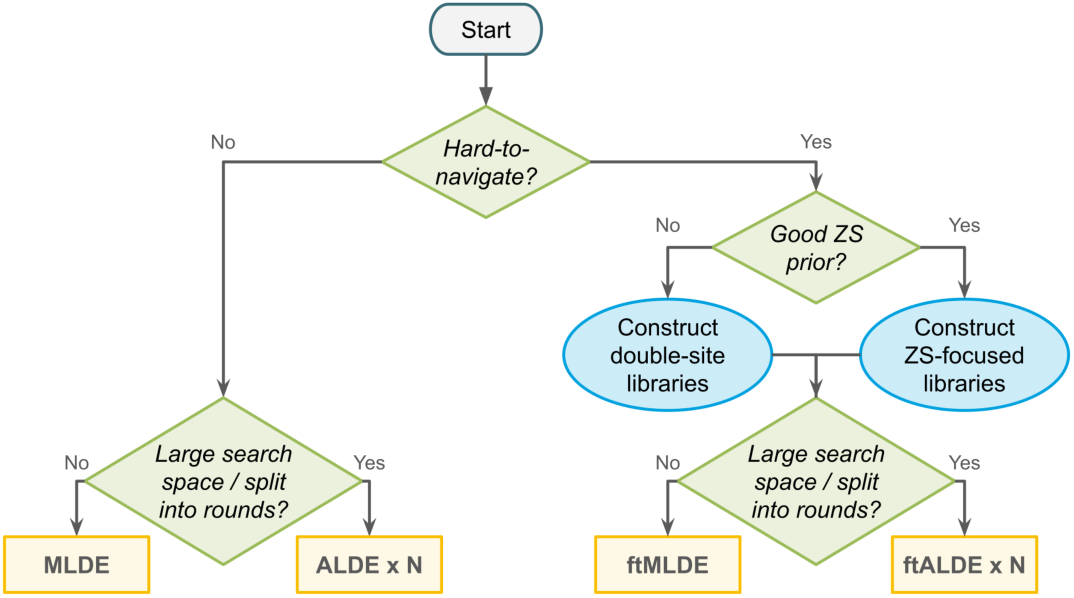
Decision tree summarizing recommended ML strategies based on total number of variants screened experimentally, landscape navigability (e.g. active variant percentage, pairwise epistasis), the quality of ZS active/inactive variant classification (i.e. ROC-AUC > 0.5), and the number of available screening rounds (N).

We focused this study on combinatorial landscapes typically generated using SSM, which are often enriched in epistasis and present challenges for DE. Our decision leveraged the observation that random mutations spread across a protein generally exhibit little sign epistasis, and thus beneficial mutations can often be combined to great success using laboratory methods such as staggered extension process (StEP) recombination to generate variants with higher fitness.^68,69^ In this context, Hamming distance has been demonstrated to have a weak correlation with variant fitness.^23^ The ZS predictor benchmark has also been performed predominantly on datasets with random mutation spread across a protein.^25^

We also evaluated several additional design choices for ML strategies that had a more limited impact on MLDE performance. First, we explored more informative ways to represent protein sequences compared to a categorical encoding (one-hot, which has no learned information and treats all amino acids equally). Learned representations from PLMs (e.g., ESM2) showed minimal to no improvement over one-hot encoding for landscapes with at least 1% active variants (Figure S21), including in the focused training setting (Figure S22). However, they did exhibit improvements for landscapes with fewer than 1% active variants (Figure S23). While not as beneficial as focused training (Figure S8), learned representations may still enhance performance for particularly challenging landscapes when combined with focused training (Figure S24). Additionally, we used boosting models to facilitate a direct comparison between MLDE and ALDE. Different model choices and ensembles, such as ridge regression for MLDE or deep neural network ensembles for ALDE, could offer further improvements (Figure S25 and S26). We also provide a codebase, SSMuLA (Site-Saturation Mutagenesis Landscape Assistant), which includes options for these granular design choices.

While we streamlined focused training design choices, we also identified areas for improvement. Based on testing individual ZS predictors across different thresholds of the original search space, we set the focused training library threshold to the top 12.5%, ranked by ZS scores, across all landscapes (Figure S27). We then naively ensembled ZS predictors to demonstrate the benefits of combining orthogonal priors (Methods). The current Hamming distance ensembled focused training libraries inherently had a size cutoff (i.e., 12.5% of a double-site library on a four-site landscape is 300). More sophisticated approaches, such as MODIFY,^70^ which balance ZS selection with training set diversity and manage the exploration-exploitation trade-off, may offer a more comprehensive and autonomous method for selecting and ensembling ZS predictors.

There are also signs that ZS predictors are intrinsically limited for certain prediction tasks. For instance, the K227 substitution in TrpB, which enables high-fitness variants under engineering conditions but is nearly undetectable in natural sequences, might not be adequately captured by EVmutation.^44^ Additionally, the performance of different ZS predictors varied even within the same protein, as observed in TrpB across different landscapes. This indicates that while natural evolutionary information can be predictive, it may fall short in capturing evolution in the laboratory. Furthermore, all the enzyme systems we studied primarily involved native or near-native functions, with a majority being TrpB landscapes. When applied to non-native functions, we expect that the usefulness of evolutionary-based ZS predictors will decrease, perhaps significantly so. In the case of the TEV and T7 landscapes, none of the ZS predictors consistently improved focused training performance from random sampling, suggesting room for the development of new ZS predictors.

In summary, our study lays the groundwork for a ML-assisted protein engineering framework leveraging the strengths of multiple ML approaches, including MLDE, ftMLDE, ALDE, and, introduced here, ftALDE. With our growing ability to read^71,72^ and write sequences,^73^ along with improved tools for constructing libraries,^74^ we believe our findings will demystify the application of ML-based DE strategies and encourage their broader adoption in protein engineering.

## Acknowledgments

The authors thank Sabine Brinkmann-Chen, Tanvi Ganapathy, Ariane Mora, Chenghao Liu, Yueming Long, Julia Reisenbauer, Casey Ritts, Kathleen Sicinski, Bruce Wittmann, Kevin Yang, and the other members of the Arnold Lab for critical reading and discussion of the manuscript, Andrei Papkou, Vikram Sundar, and Boqiang Tu for dataset discussion, and Thomas Hopf and Aviv Spinner for assistance with EVmutation implementation.

This work was supported by the NSF Division of Chemical, Bioengineering, Environmental and Transport Systems (CBET 1937902) and Amgen Chem-Bio-Engineering Award (CBEA AMGEN.ARNOLD22). F.Z.L. and J.Y. were partially supported by the National Science Foundation Graduate Research Fellowship and F.Z.L. was partially supported by Amazon AI4Science Fellowship at Caltech.

## Author contributions

F.Z.L., conceptualization, data curation, formal analysis, investigation, methodology, software, visualization, writing – original draft, writing – review & editing

J.Y., conceptualization, methodology, software, writing -review & editing K.E.J., conceptualization, methodology, software, writing – review & editing

E. G., software

Y. Y., funding acquisition, resources, supervision, writing – review & editing F.H.A., funding acquisition, resources, supervision, writing – review & editing

## Declaration of generative AI and AI-assisted technologies in the writing process

During the preparation of this work the authors used ChatGPT in order to check spelling, grammar, and improve the readability and language of the manuscript. After using this tool, the authors reviewed and edited the content as needed and take full responsibility for the content of the published article.

## Resource availability

### Lead contact

Further information and requests for resources should be directed to and will be fulfilled by the lead contact, Frances Arnold.

### Materials availability

Not applicable to this study.

### Data and code availability

All data and results that support this study are deposited at https://doi.org/10.5281/zenodo.13910506. All code is available at https://github.com/fhalab/SSMuLA.

## Methods

| REAGENT or RESOURCE | SOURCE | IDENTIFIER |
| --- | --- | --- |
| Deposited data |  |  |
| Bacterial toxin-antitoxin (ParD-ParE) fitness landscape | Lite, *et al.*, 2020 | https://doi.org/10.7554/eLife.60924 |
| Bacterial toxin-antitoxin (ParD-ParE) structure | Protein Data Bank | PDB: 6X0A; PDB: 5CEG |
| Protein G B1 domain (GB1) fitness landscape | Wu, *et al.*, 2016 | https://doi.org/10.7554/eLife.16965 |
| Protein G B1 domain (GB1) structure | Protein Data Bank | PDB: 2GI9 |
| Dihydrofolate reductase (DHFR) fitness landscape | Papkou, *et al.*, 2023 | https://doi.org/10.1126/science.adh3860 |
| Dihydrofolate reductase (DHFR) structure | Protein Data Bank | PDB: 6XG5 |
| T7 RNA polymerase fitness landscape | Tu, *et al.*, 2024 | https://doi.org/10.1101/2022.03.09.483646 |
| T7 RNA polymerase structure | Protein Data Bank | PDB: 1CEZ |
| TEV protease fitness landscape | Tu, *et al.*, 2024 | https://doi.org/10.1101/2022.03.09.483646 |
| TEV protease structure | Protein Data Bank | PDB: 1LVM |
| β-subunit of tryptophan synthase (TrpB) fitness landscape | Johnston, *et al.*, 2024 | https://doi.org/10.1073/pnas.2400439121 |
| β-subunit of tryptophan synthase (TrpB) structure | Protein Data Bank | PDB: 8VHH |
| Compiled data and results | This paper | 10.5281/zenodo.13910506 |
| Software and algorithms |  |  |
| SSMuLA code and anaconda environment | This paper | https://github.com/fhalab/SSMuLA |
| EVmutation | Hopf, *et al.*, 2017 | https://v2.evcouplings.org/ |
| Evolutionary Scale Modeling (ESM) | Rives, *et al.*, 2021;<br>Meier, *et al.*, 2021;<br>Lin, *et al.*, 2023 | https://github.com/facebookresearch/esm |
| ESM inverse folding (ESM-IF) | Hsu, *et al.*, 2022 | https://github.com/facebookresearch/esm |
| Combinatorial Variant Effects from Structure (CoVES) | Ding, *et al.*, 2024 | https://github.com/ddingding/CoVES |
| Triad | Protabit, Pasadena, CA, USA | https://triad.protabit.com/ |
| Machine Learning-Assisted Directed Evolution (MLDE) | Wittmann, *et al.*, 2021 | https://github.com/fhalab/MLDE |
| Active Learning-Assisted Directed Evolution | Yang, *et al.*, 2024 | https://github.com/jsunn-y/ALDE |

### Landscape preparation

By choosing essentially complete datasets, we minimized the need for data imputation, thus avoiding potential biases and misrepresentations. We focused on libraries with at least 1% active variants in the main text to keep the main text relevant to the majority of campaigns, with results for all landscapes provided in the supplementary material. All fitness values were normalized so that the variant with the maximum fitness has a value of one.

### Landscape attributes

We considered two groups of attributes for this analysis: 1) fitness statistics, which included percent of active variants and parameters derived from simple statistical modeling, and 2) ruggedness, which included pairwise epistasis and the number of local optima. We do not impute missing values.

#### Definition of active variants

For landscapes containing fitness data for variants with stop codons, “active” variants were defined as those 1.96 standard deviations above the mean fitness of all sequences containing stop codons, which are expected to be inactive.^44^ For GB1, T7, and TEV we followed the cutoffs set by the authors, based on the detection limit of their fitness measurement system.^37,42,43^

#### Fitness statistics

We used the “statistical functions” (’scipy.stats’) and signal (’scipy.signal’) modules from the SciPy Python package^75^ to calculate kurtosis, estimate the Cauchy peak location, and determine the number of KDE peaks. Specifically, kurtosis was calculated using the ’kurtosis’ function with default settings from the ’stats’ module. Cauchy peak location was estimated using the ’fit’ method from the ’cauchy’ distribution object in the ’stats’ module. The number of KDE peaks was determined by estimating the probability density function with the ’gaussian_kdè function from the ’stats’ module and then identifying local optima using the ’argrelextremà function from the signal module.

#### Pairwise epistasis calculation

We classified pairwise epistasis into three categories: magnitude, sign, and reciprocal sign. For each active variant, we assigned an epistasis type for each possible double substitution at chosen sites.^44^ We then calculated the fraction of epistasis type for each starting variant in the landscape. To enhance relevance to DE navigability, we incorporated additive interactions into magnitude epistasis, and merged sign and reciprocal sign epistasis into non-magnitude epistasis. Missing values are omitted.

##### Magnitude epistasis

The combined effect of two mutations is larger than or equal to their additive effects in the same direction as each individual mutation. This is navigable through single-step or recombination-based DE methods.

##### Sign epistasis

The direction of the effect of one mutation changes in the presence of the other such that the substitution order impacts single-step DE navigability.

##### Reciprocal sign

The combined effect changes the direction of both mutations in the presence of each other. This is not accessible with single-step DE that is inherently a greedy uphill walk.

#### Pairwise epistasis correlation with C-alpha distances

The pairwise C-alpha distances of mutated residues were calculated based on each of the parent structure and then averaged for each landscape. The Spearman’s correlation was then calculated with fraction of pairwise non-magnitude epistasis

#### Local optimum calculation

A local optimum is a variant with higher fitness than all its neighboring active variants differing by one amino acid substitution.^41,44,46,47,49^

### ZS calculation

We calculated six different ZS scores for each landscape. The ranking correlation was calculated using Spearman’s correlation and the active/inactive variant classification was quantified by ROC-AUC. All values can be found with data deposit.

#### Hamming distance

Hamming distance counted the number of amino acid differences between a variant and the parent sequence. In the main text, the parent sequence was defined by the authors of each landscape. Simulations provided in the supplemental information explored all possible parent sequences, starting from any active variant. The Hamming distance from each given parent sequence was used for fitness ranking and active/inactive variant classification, with the final results averaged over all possible parent sequences.

#### EVmutation score

EVmutation score was based on conservation and evolutionary couplings with multiple sequence alignments (MSAs).^18,56^ For each landscape, the parent sequence was uploaded to the EVcoupling web server with default parameters for MSA generation and their subsequent EVmutation model training. We chose the recommended EVmutation model if the alignment covers all mutation sites, and otherwise we prioritized the models covering all sites with a higher bitscore. In the case that not all the mutation sites are covered, the position filter was decreased from the default 70% to 50%. The EVmutation scores and ranking were then generated with the EVmutation model.

#### ESM score

ESM score was based on pretrained protein language model masked language modeling objective output probability for mutations given their surrounding context.^19^ Contrary to EVmutation, ESM score does not entail explicit MSAs. For each landscape, ESM score for each position was calculated by using the log odds ratio comparing the mutated amino acid probability with the parent probability. Multi-mutant ESM scores are then summed from individual mutants. We found ESM-1v (esm1v_t33_650M_UR90S_1), ESM-1b (esm1b_t33_650M_UR50S), and ESM2 (esm2_t33_650M_UR50D) giving comparable results and decided to move forward with ESM2.

#### ESM-IF score

ESM-IF score was calculated using the ratio between likelihoods of the mutated and parent sequences according to the inverse folding model ESM-IF1 (esm_if1_gvp4_t16_142M_UR50),^20^ given the experimentally determined parent structure from PDB.^76^ It leverages a Geometric Vector Perceptron (GVP) module that maintains properties under transformations like rotations.^77^

#### CoVES score

The CoVES score was calculated using pretrained weights from Ding *et al.* (2024) following their methods,^21^ applied to all parent structures from the PDB.

#### Triad score

Triad score reflected the change in free energy of folding upon mutation (ΔΔG) prediction of mutant stability. The calculation was based on Rosetta energy functions across mutation sites with a fixed backbone assumption. First, the parent protein crystal structure for each landscape was obtained from PDB. Then, for each landscape, we obtained the score for each variant with the Triad software suite following the method from Wittmann *et al.* (2021).^12^

### ZS ensembles

For each landscape, the six different ZS scores were calculated accordingly. For Hamming distance-based ensemble, a Hamming distance of two was first applied (i.e., double-site libraries), then five other ZS scores were used to rank the variants. For other ensembles, equal weight was given to each individually calculated and ranked ZS score for each variant.

### ZS analysis

#### ZS MSA depth correlation

The MSA depth referred to the number of sequences resulted from EVmutation, where all mutation sites were covered.

#### ZS pairwise correlation

The pairwise correlation was performed for each landscape and then averaged across the 12 landscapes with at least 1% active variants.

### DE simulations

For each landscape, all DE simulations started from an active variant, regardless of its background. The maximum fitness achieved by each starting variant was recorded.

#### Single-step DE

This is a greedy walk algorithm. The process begins with selecting one of the possible substitution sites, evaluating the fitness impact of all possible amino acid substitutions at this position. The substitution yielding the highest fitness is fixed, and the position is restricted from further exploration. In the next round, one of the remaining positions is selected, with all mutants evaluated, and the best substitution is fixed again. This process repeats iteratively until all positions have been evaluated yielding the fitness of the best variant identified in the last round. Consequently, each site is optimized once per simulation. For example, a four-site library requires four rounds of single-step DE to reach the optimal variant and there is a total of 24 (4!) possible orders of sampling. This is a deterministic approach to navigate the fitness landscape as the best variant is always selected.^11,12,44^

#### Recombination SSM

This is a naive recombination. This approach randomly samples the combinatorial space, independently optimizing each site within the context of the initial sequence and then combining the best substitutions from each site into a new variant.^11,44^

#### Top96 recombination

This is an alternative recombination approach. All substitutions are made at each of the sites independently in the background of the initial sequence, calculating fitness for all combinations from single substitution over the initial sequence. The sequences are then ranked based on their fitness, and the top 96 variants are tested *in silico*. The reported maximum fitness reflects the highest observed among the initial sequence, any single substitutions, and the best of the top 96.^44^

### MLDE, ALDE, and focus-training experiments

For each experiment on a given landscape, a range of total number of samples (sizes: 120, 144, 192, 288, 384, 480, 576, 672, 1056, and 2016) were split across training and testing for MLDE or multiple rounds of sampling for ALDE. All results were averaged across 50 replicates. The top 96 predicted variant fitness values were analyzed.

#### Encoding strategies

One-hot and learned representations from ESM2 (esm2_t33_650M_UR50D) were tested. One-hot encodings were flattened over the mutated sites. Learned representations from ESM2 (esm2_t33_650M_UR50D) were implemented in three ways, (1) flattened over the mutated sites, (2) mean pooled over the mutated sites, and (3) mean pooled over the full sequence.

#### MLDE experiments

For each MLDE experiment on a given landscape, XGBoost^78^ and the Scikit-learn ridge regression^79^ models were trained on different random samples (sizes: 24, 48, 96, 192, 288, 384, 480, 576, 960, and 1920) with five-fold cross-validation. An alpha value of 1 was used for ridge regression. The ’reg:tweediè objective was implemented with an ’early_stopping_rounds’ of 10 for the boosting models. The model ensembles were used to predict variant fitness across the entire library, and the top 96 predicted variant fitness values were analyzed.

#### ALDE experiments

For each ALDE experiment on a given landscape, models were trained on different random samples (total number: 120, 144, 192, 288, 384, 480, 576, 672, 1056, and 2016) split across different iterations (rounds: 2, 3, and 4). Boosting ensemble and deep neural network ensembles were tested for ALDE. The ’reg:tweediè objective was implemented with an ’early_stopping_rounds’ of 10 for the boosting models. The ’torch.optim.Adam’ optimizer with the ’torch.nn.MSELoss’ loss from PyTorch^80^ was implemented for deep neural network ensembles with bootstrapping of five models where 90% of the total training data was randomly seen during training. Greedy acquisition functions were deployed for both boosting and deep neural network ensembles.

#### Focused training experiments

Different focused training sets (50%, 25%, and 12.5% of total mutants) were ranked by favorable ZS scores with the chosen ZS predictor for each given landscape. These focused sets were then used to sample training data for MLDE or the initial round of ALDE. 12.5% was chosen for all simulations other than examining the optimal focused training library size (Figure S27).

### Feature correlation and importance analysis

To analyze how each landscape attribute correlated with the simulation targets, a Spearman’s correlation was performed between the attribute and the performance of the model. Both the Spearman’s ρ and p-value were reported. To test the differences between binding and enzyme activities, t-tests were performed, where the t-statistic and p-values were reported. A p-value less than 0.05 was considered statistically significant.

## Supplemental information

**Table S1.**
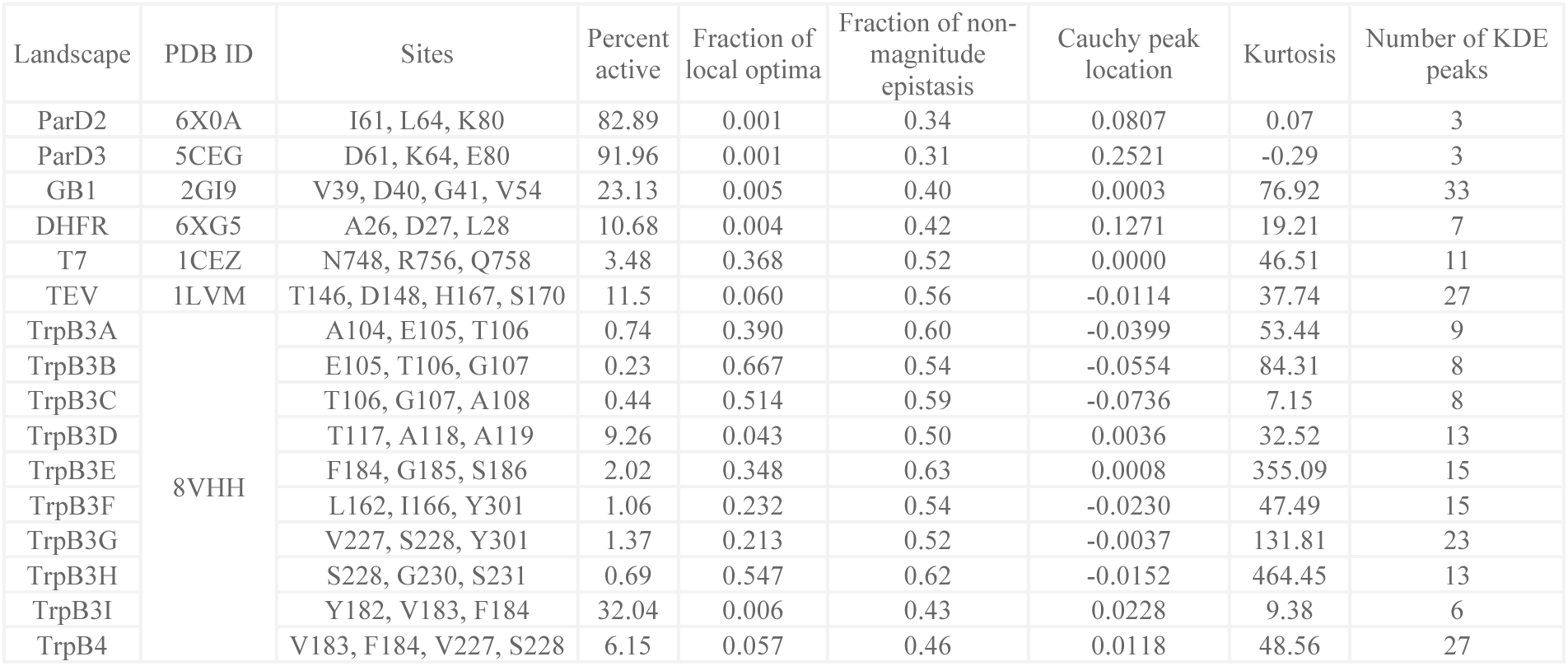
Combinatorial landscapes with additional details including landscapes with fewer than 1% active variants, related to Table 1.^37,40–44^.

| Landscape | PDB ID | Sites | Percent active | Fraction of local optima | Fraction of non-magnitude epistasis | Cauchy peak location | Kurtosis | Number of KDE peaks |
| --- | --- | --- | --- | --- | --- | --- | --- | --- |
| ParD2 | 6X0A | I61, L64, K80 | 82.89 | 0.001 | 0.34 | 0.0807 | 0.07 | 3 |
| ParD3 | 5CEG | D61, K64, E80 | 91.96 | 0.001 | 0.31 | 0.2521 | -0.29 | 3 |
| GB1 | 2GI9 | V39, D40, G41, V54 | 23.13 | 0.005 | 0.40 | 0.0003 | 76.92 | 33 |
| DHFR | 6XG5 | A26, D27, L28 | 10.68 | 0.004 | 0.42 | 0.1271 | 19.21 | 7 |
| T7 | 1CEZ | N748, R756, Q758 | 3.48 | 0.368 | 0.52 | 0.0000 | 46.51 | 11 |
| TEV | 1LVM | T146, D148, H167, S170 | 11.5 | 0.060 | 0.56 | -0.0114 | 37.74 | 27 |
| TrpB3A | 8VHH | A104, E105, T106 | 0.74 | 0.390 | 0.60 | -0.0399 | 53.44 | 9 |
| TrpB3B |  | E105, T106, G107 | 0.23 | 0.667 | 0.54 | -0.0554 | 84.31 | 8 |
| TrpB3C |  | T106, G107, A108 | 0.44 | 0.514 | 0.59 | -0.0736 | 7.15 | 8 |
| TrpB3D |  | T117, A118, A119 | 9.26 | 0.043 | 0.50 | 0.0036 | 32.52 | 13 |
| TrpB3E |  | F184, G185, S186 | 2.02 | 0.348 | 0.63 | 0.0008 | 355.09 | 15 |
| TrpB3F |  | L162, I166, Y301 | 1.06 | 0.232 | 0.54 | -0.0230 | 47.49 | 15 |
| TrpB3G |  | V227, S228, Y301 | 1.37 | 0.213 | 0.52 | -0.0037 | 131.81 | 23 |
| TrpB3H |  | S228, G230, S231 | 0.69 | 0.547 | 0.62 | -0.0152 | 464.45 | 13 |
| TrpB3I |  | Y182, V183, F184 | 32.04 | 0.006 | 0.43 | 0.0228 | 9.38 | 6 |
| TrpB4 |  | V183, F184, V227, S228 | 6.15 | 0.057 | 0.46 | 0.0118 | 48.56 | 27 |

**Table S2.** MLDE percent improvement from three types of DE, related to Figure 2a.

| Number of MLDE training samples | Recomb |  | Single-step |  | Top96 recomb |  |
| --- | --- | --- | --- | --- | --- | --- |
|  | Average max fitness achieved | Fraction reaching the global optimum | Average max fitness achieved | Fraction reaching the global optimum | Average max fitness achieved | Fraction reaching the global optimum |
| 24 | -7.35 | -31.60 | -15.25 | -67.40 | -23.89 | -79.13 |
| 48 | 3.13 | 30.59 | -5.66 | -37.76 | -15.28 | -60.16 |
| 96 | 18.29 | 108.32 | 8.20 | -0.71 | -2.84 | -36.44 |
| 192 | 28.66 | 162.73 | 17.69 | 25.22 | 5.68 | -19.84 |
| 288 | 33.27 | 201.60 | 21.91 | 43.75 | 9.47 | -7.98 |
| 384 | 34.95 | 238.91 | 23.44 | 61.53 | 10.85 | 3.40 |
| 480 | 37.94 | 282.44 | 26.17 | 82.28 | 13.30 | 16.69 |
| 576 | 40.24 | 313.53 | 28.28 | 97.09 | 15.20 | 26.17 |
| 960 | 47.09 | 427.02 | 34.54 | 151.18 | 20.82 | 60.80 |
| 1920 | 50.85 | 487.65 | 37.99 | 180.08 | 23.92 | 79.30 |

**Table S3.** ftMLDE percent improvement from MLDE, related to Figure 2a.

| Number of MLDE training samples | Average max fitness achieved | Fraction reaching the global optimum |
| --- | --- | --- |
| 24 | 11.17 | 76.89 |
| 48 | 11.86 | 53.57 |
| 96 | 9.57 | 60.32 |
| 192 | 8.64 | 76.73 |
| 288 | 7.08 | 70.02 |
| 384 | 6.94 | 58.94 |
| 480 | 5.18 | 44.44 |
| 576 | 4.04 | 38.85 |
| 960 | 0.13 | 17.60 |
| 1920 | -1.60 | 9.48 |

**Table S4.** MLDE, ALDE and focused training with 480 total sample size fold improvement from single-step DE, related to Figure 2c. Bold row indicates the landscape with the max improvement and italic rows indicate landscapes with fewer than 1% active variants.

| Landscape | MLDE | ftMLDE | ALDE | ftALDE | ALDE x 3 | ftALDE x 3 | ALDE x 4 | ftALDE x 4 |
| --- | --- | --- | --- | --- | --- | --- | --- | --- |
| DHFR | 1.08 | 1.07 | 1.04 | 1.07 | 1.06 | 1.08 | 1.05 | 1.07 |
| GB1 | 1.17 | 1.34 | 1.16 | 1.27 | 1.25 | 1.37 | 1.34 | 1.38 |
| ParD2 | 1.04 | 1.04 | 1.04 | 1.04 | 1.04 | 1.04 | 1.04 | 1.04 |
| ParD3 | 1.01 | 1.01 | 1.02 | 1.02 | 1.02 | 1.02 | 1.02 | 1.02 |
| T7 | 1.40 | 1.32 | 1.31 | 1.31 | 1.38 | 1.34 | 1.38 | 1.33 |
| TEV | 1.42 | 1.28 | 1.28 | 1.26 | 1.38 | 1.32 | 1.42 | 1.31 |
| *TrpB3A* | *1.52* | *2.20* | *1.31* | *1.84* | *1.60* | *2.02* | *1.61* | *2.15* |
| *TrpB3B* | *1.25* | *2.05* | *0.85* | *1.47* | *0.96* | *1.44* | *0.87* | *1.48* |
| *TrpB3C* | *1.09* | *1.71* | *0.89* | *1.43* | *0.88* | *1.35* | *0.98* | *1.37* |
| TrpB3D | 1.24 | 1.32 | 1.26 | 1.28 | 1.30 | 1.30 | 1.30 | 1.32 |
| **TrpB3E** | **2.30** | **3.48** | **1.43** | **2.84** | **1.97** | **2.96** | **1.71** | **2.96** |
| TrpB3F | 1.30 | 1.43 | 1.38 | 1.39 | 1.40 | 1.43 | 1.37 | 1.43 |
| TrpB3G | 1.38 | 1.55 | 1.30 | 1.49 | 1.37 | 1.52 | 1.49 | 1.55 |
| *TrpB3H* | *1.30* | *2.46* | *0.89* | *2.10* | *1.07* | *2.13* | *1.52* | *2.28* |
| TrpB3I | 1.21 | 1.26 | 1.24 | 1.25 | 1.25 | 1.26 | 1.25 | 1.26 |
| TrpB4 | 1.24 | 1.34 | 1.10 | 1.25 | 1.31 | 1.35 | 1.36 | 1.41 |

**Table S5.** Protein function and MSA impact ZS predictor performances test significance, related to Figure 3c. Correlation between Spearman’s correlation of ZS predictor fitness ranking prediction with MSA depth, where the depth for the EVmutation calculation covering the full sequence is used. Bold font indicates statistically significant (p-value < 0.05).

| Metric | MSA depth<br>(Spearman's correlation) |  |
| --- | --- | --- |
| | Spearman $\rho$ | p-value |
| Hamming distance | 0.54 | 0.07 |
| EVmutation | 0.49 | 0.11 |
| ESM | 0.71 | **0.01** |
| ESM-IF | 0.55 | 0.07 |
| CoVES | 0.03 | 0.93 |
| Triad | -0.05 | 0.87 |

**Table S6.** T-test for ZS predictor between binding and enzyme activities for landscapes with at least 1% active variants, related to Figure 4a. Bold font indicates statistically significant (p-value < 0.05).

| Metric | Binding vs. Enzyme activities<br>(Spearman's correlation) |  | Binding vs. Enzyme activities<br>(ROC-AUC) |  |
| --- | --- | --- | --- | --- |
|  | t-statistics | p-value | t-statistics | p-value |
| Hamming distance | 1.740 | 0.210 | -2.379 | **0.042** |
| EVmutation | 1.738 | 0.167 | -1.669 | 0.126 |
| ESM | 1.297 | 0.308 | -0.747 | 0.493 |
| ESM-IF | 3.316 | 0.052 | 0.749 | 0.494 |
| CoVES | 3.641 | 0.057 | 1.289 | 0.279 |
| Triad | 4.332 | **0.001** | 1.101 | 0.334 |

**Table S7.** T-test for focused training MLDE (480 total sample size) between binding and enzyme activities for landscapes with at least 1% active variants, related to Figure 4b.

| Metric | Binding vs. Enzyme activities<br>(Spearman's correlation) |  | Binding vs. Enzyme activities<br>(ROC-AUC) |  |
| --- | --- | --- | --- | --- |
|  | t-statistics | p-value | t-statistics | p-value |
| ZS predictor |  |  |  |  |
| Hamming distance | 0.041 | 0.969 | -0.720 | 0.525 |
| EVmutation | 0.351 | 0.738 | -0.111 | 0.918 |
| ESM | 0.341 | 0.784 | -0.338 | 0.758 |
| ESM-IF | 0.582 | 0.577 | -0.267 | 0.802 |
| CoVES | 0.342 | 0.745 | 0.217 | 0.843 |
| Triad | 0.904 | 0.397 | -0.210 | 0.845 |
| Hamming distance<br>+ EVmutation | 1.445 | 0.184 | 1.313 | 0.219 |

**Table S8.**
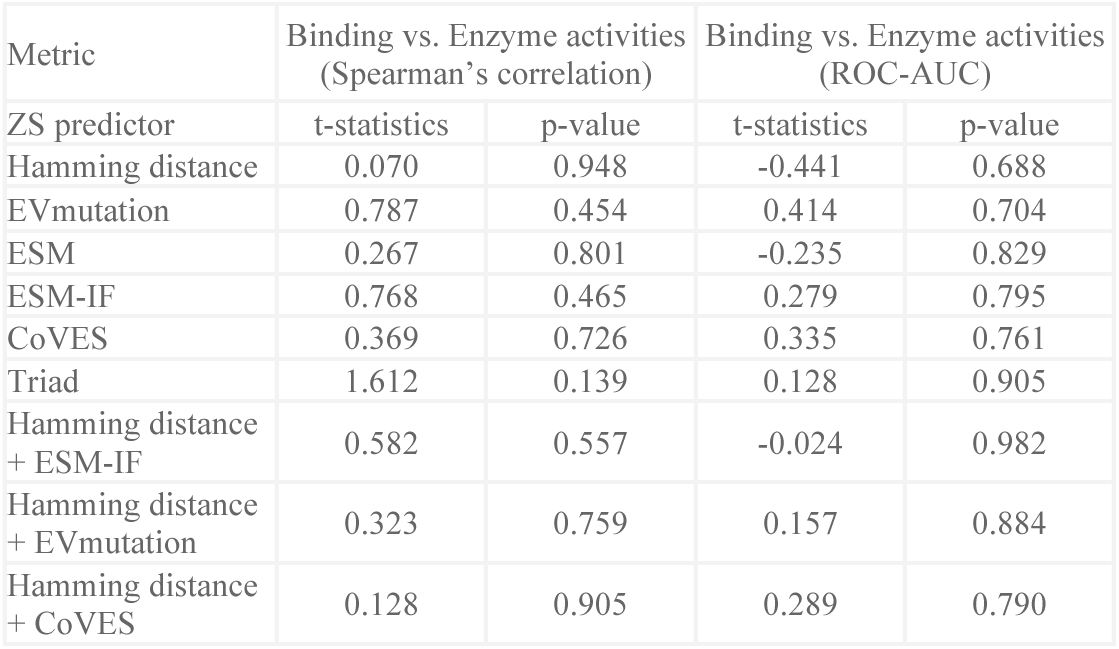
T-test for focused training ALDE (480 total sample size split into four rounds) between binding and enzyme activities for landscapes with at least 1% active variants, related to Figure 4b.

| Metric | Binding vs. Enzyme activities<br>(Spearman's correlation) |  | Binding vs. Enzyme activities<br>(ROC-AUC) |  |
| --- | --- | --- | --- | --- |
|  | t-statistics | p-value | t-statistics | p-value |
| ZS predictor |  |  |  |  |
| Hamming distance | 0.070 | 0.948 | -0.441 | 0.688 |
| EVmutation | 0.787 | 0.454 | 0.414 | 0.704 |
| ESM | 0.267 | 0.801 | -0.235 | 0.829 |
| ESM-IF | 0.768 | 0.465 | 0.279 | 0.795 |
| CoVES | 0.369 | 0.726 | 0.335 | 0.761 |
| Triad | 1.612 | 0.139 | 0.128 | 0.905 |
| Hamming distance<br>+ ESM-IF | 0.582 | 0.557 | -0.024 | 0.982 |
| Hamming distance<br>+ EVmutation | 0.323 | 0.759 | 0.157 | 0.884 |
| Hamming distance<br>+ CoVES | 0.128 | 0.905 | 0.289 | 0.790 |

**Table S9.** T-test for ZS predictor between binding and enzyme activities for landscapes with at least 1% active variants but with single substitution only, related to discussion. Bold font indicates statistically significant (p-value < 0.05).

| Metric | Binding vs. Enzyme activities<br>(Spearman's correlation) |  | Binding vs. Enzyme activities<br>(ROC-AUC) |  |
| --- | --- | --- | --- | --- |
|  | t-statistics | p-value | t-statistics | p-value |
| ZS predictor |  |  |  |  |
| Hamming distance | 0.336 | 0.746 | -3.352 | **0.010** |
| EVmutation | -0.803 | 0.478 | -0.116 | 0.924 |
| ESM | 0.467 | 0.660 | 0.521 | 0.668 |
| ESM-IF | 1.074 | 0.367 | 4.078 | **0.008** |
| CoVES | 1.032 | 0.371 | 3.116 | **0.044** |
| Triad | 1.534 | 0.205 | 5.036 | **0.001** |

**Figure S1.**
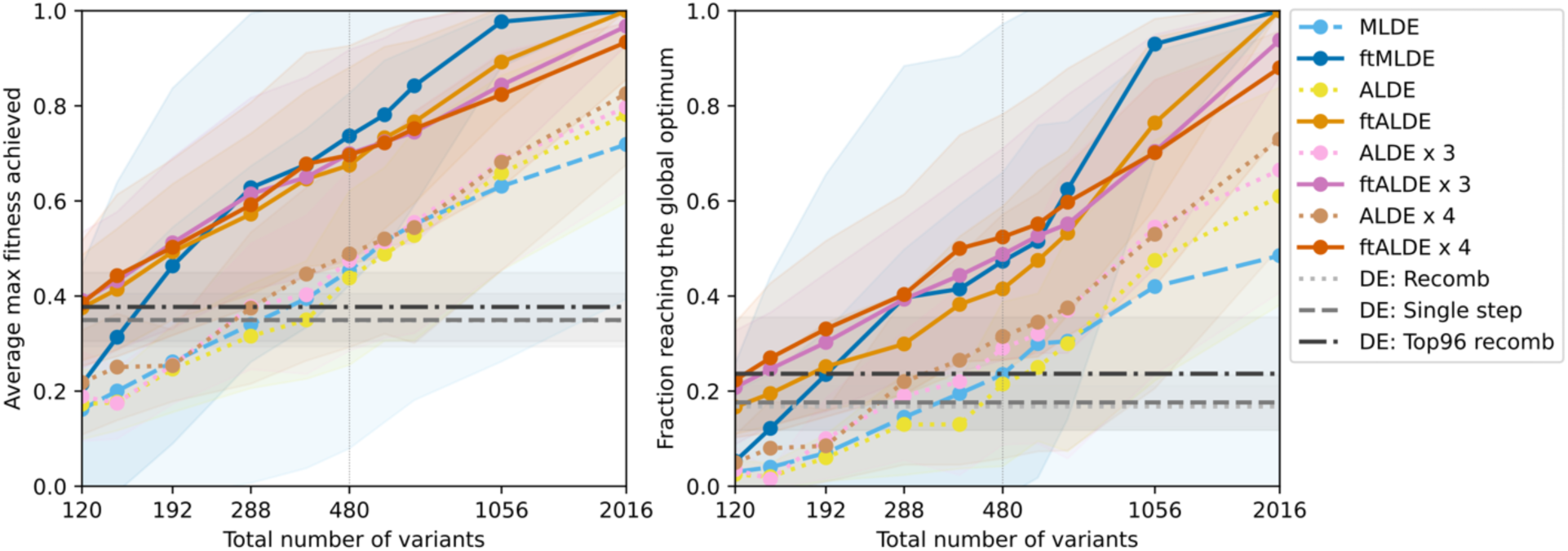
MLDE and ftMLDE performance averaged across four landscapes with fewer than 1% active variants, related to Figure 2a. Shading indicates standard deviation.

**Figure S2.**
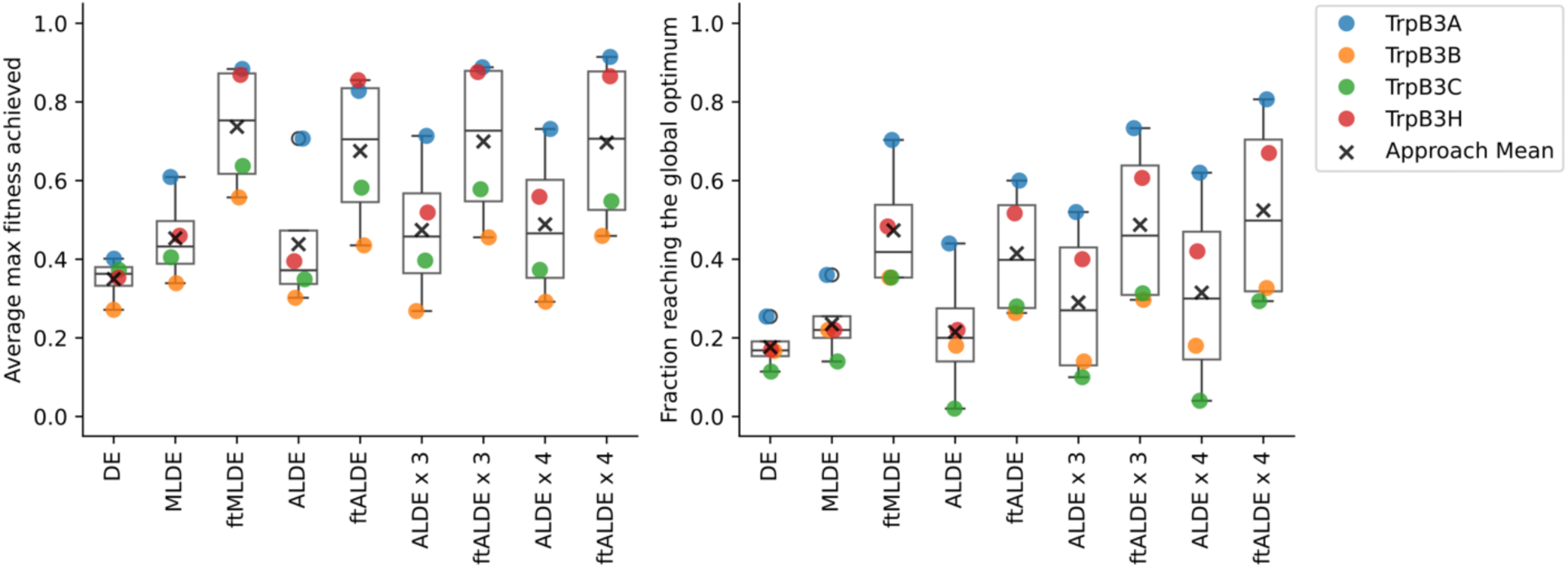
Single-step DE, MLDE, ALDE, and focused training results broken down by four landscapes with fewer than 1% active variants. A total sample size of 480 was used for all ML strategies across both metrics, related to Figure 2b.

**Figure S3.**
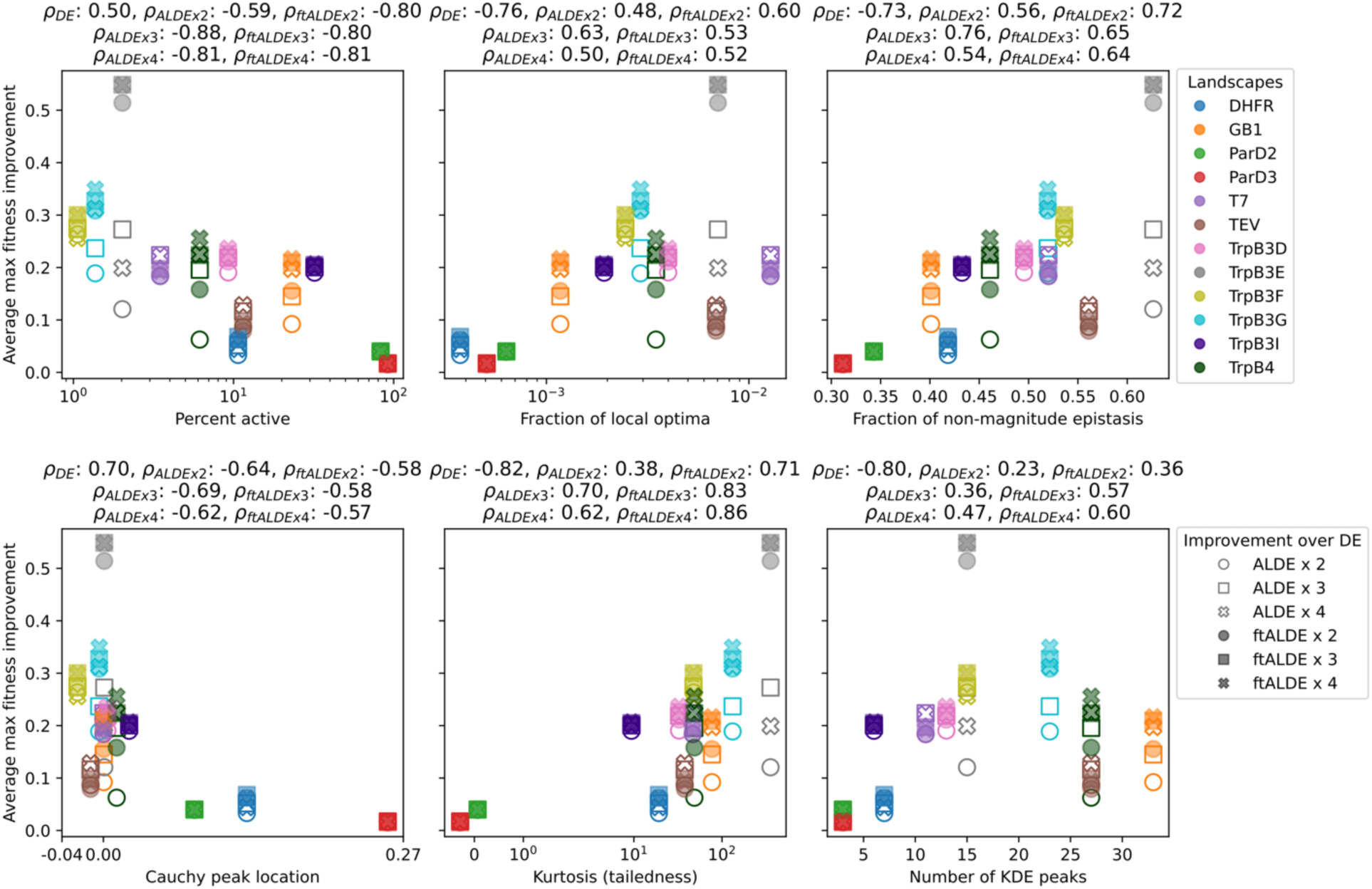
Correlation of ALDE and ftALDE performance improvement (the average maximum fitness of the top 96 predicted variants by ALDE and ftALDE over single-step DE, y-axis) with six landscape attributes (x-axis), related to Figure 2c.

**Figure S4.**
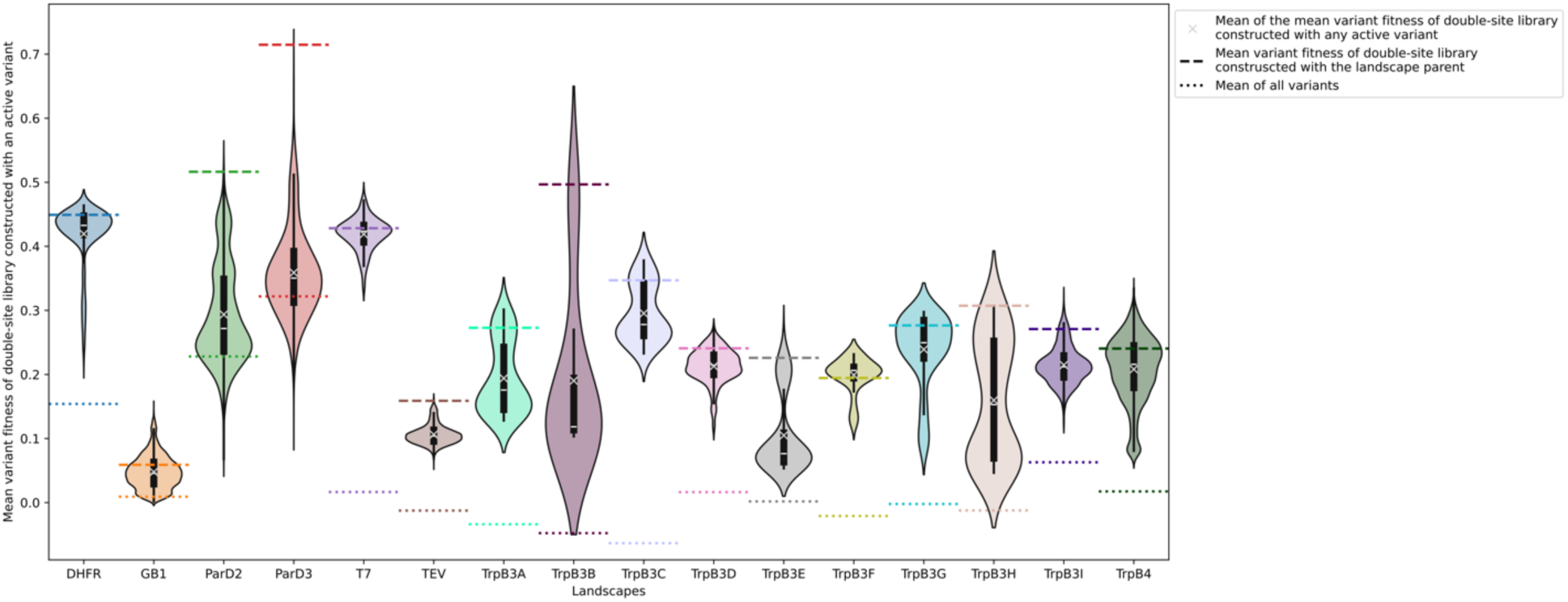
Mean variant fitness of double-site library (Hamming distance of two) from active variant as the parent, related to Hamming distance in Figure 3.

**Figure S5.**
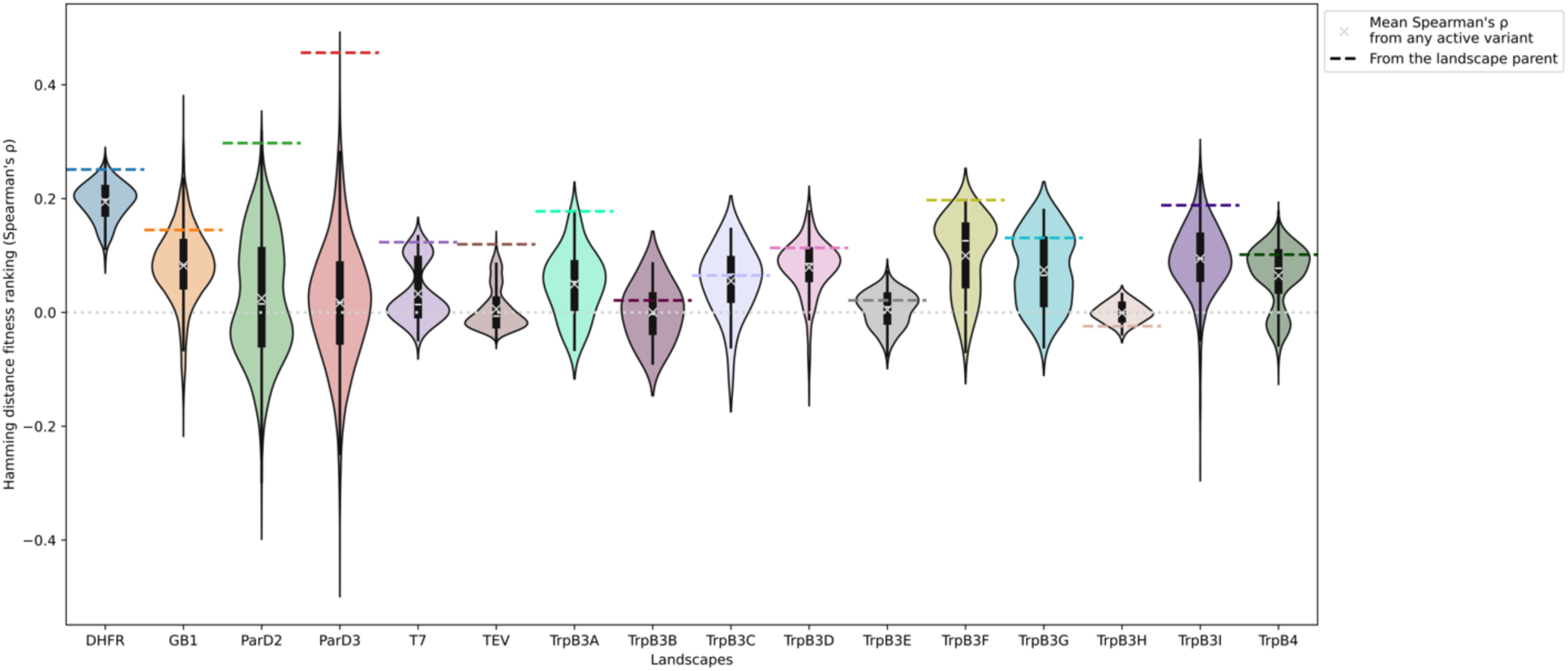
Hamming distance fitness ranking using any active variant as the parent, related to Hamming distance in Figure 3. The dotted line indicates random predictions.

**Figure S6.**
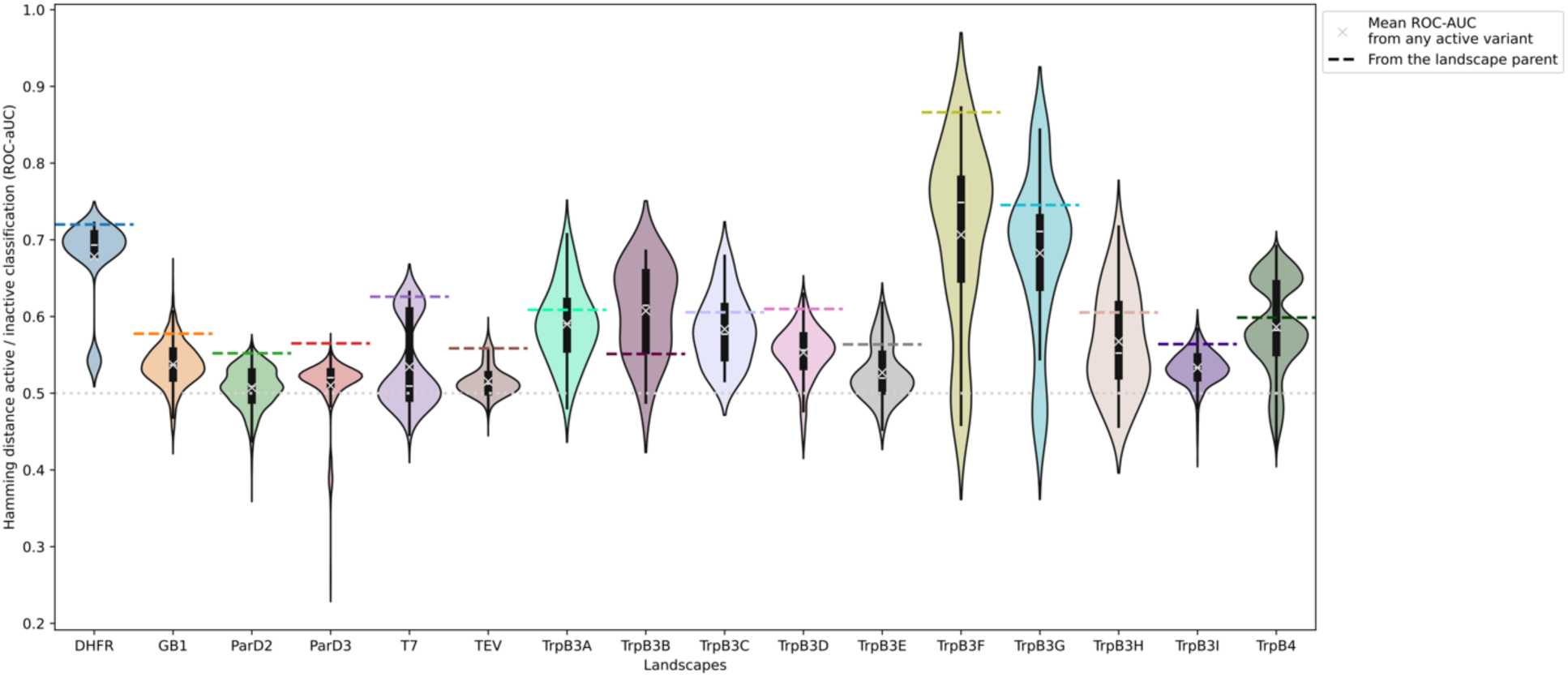
Hamming distance active/inactive variant classification using any active variant as the parent, related to Hamming distance in Figure 3. The dotted line indicates random predictions.

**Figure S7.**
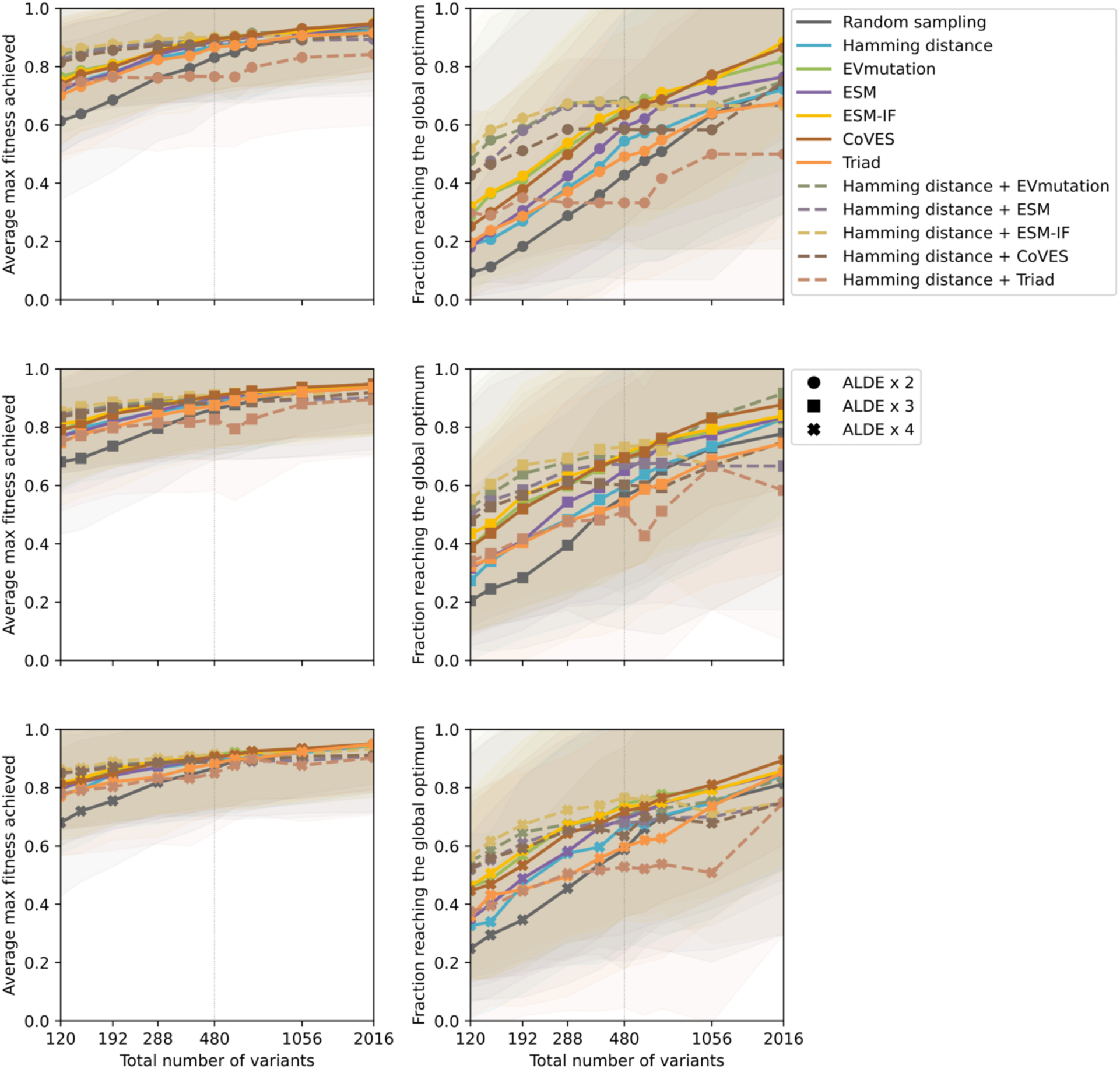
Multiple rounds of ftALDE averaged across 12 landscapes with more than 1% active variants, related to Figure 3e. Shading indicates standard deviation.

**Figure S8.**
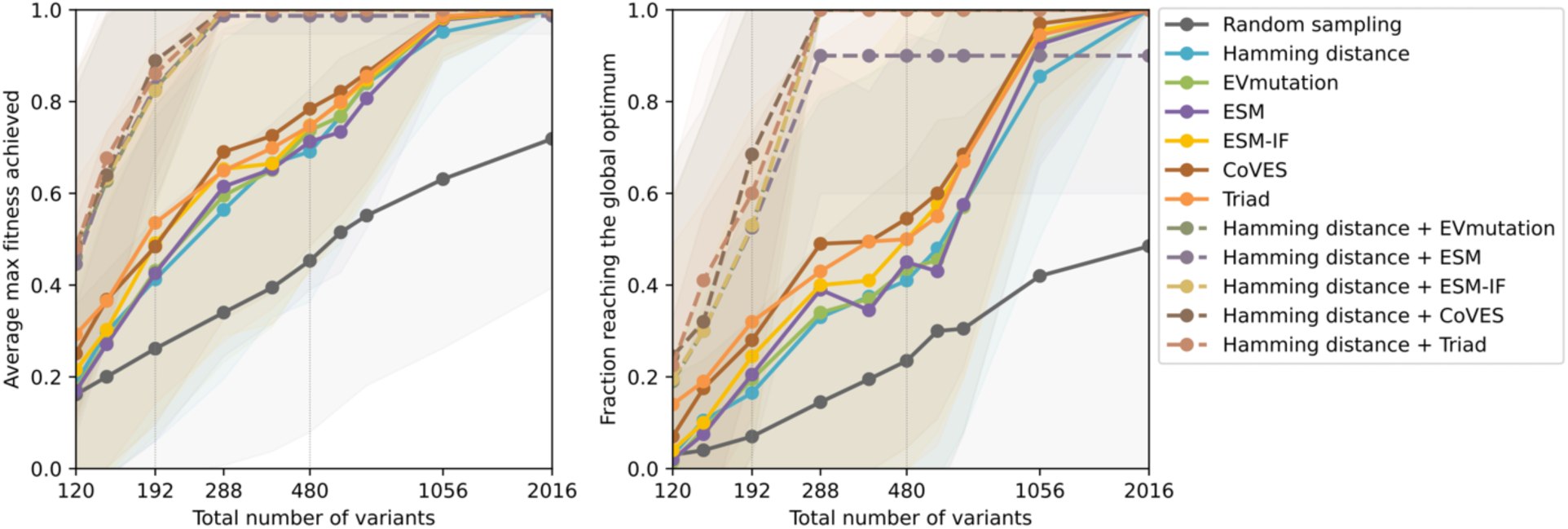
ftMLDE with Hamming distance-ensembled ZS predictors averaged across four landscapes with fewer than 1% active variants, related to Figure 3e. Shading indicates standard deviation.

**Figure S9.**
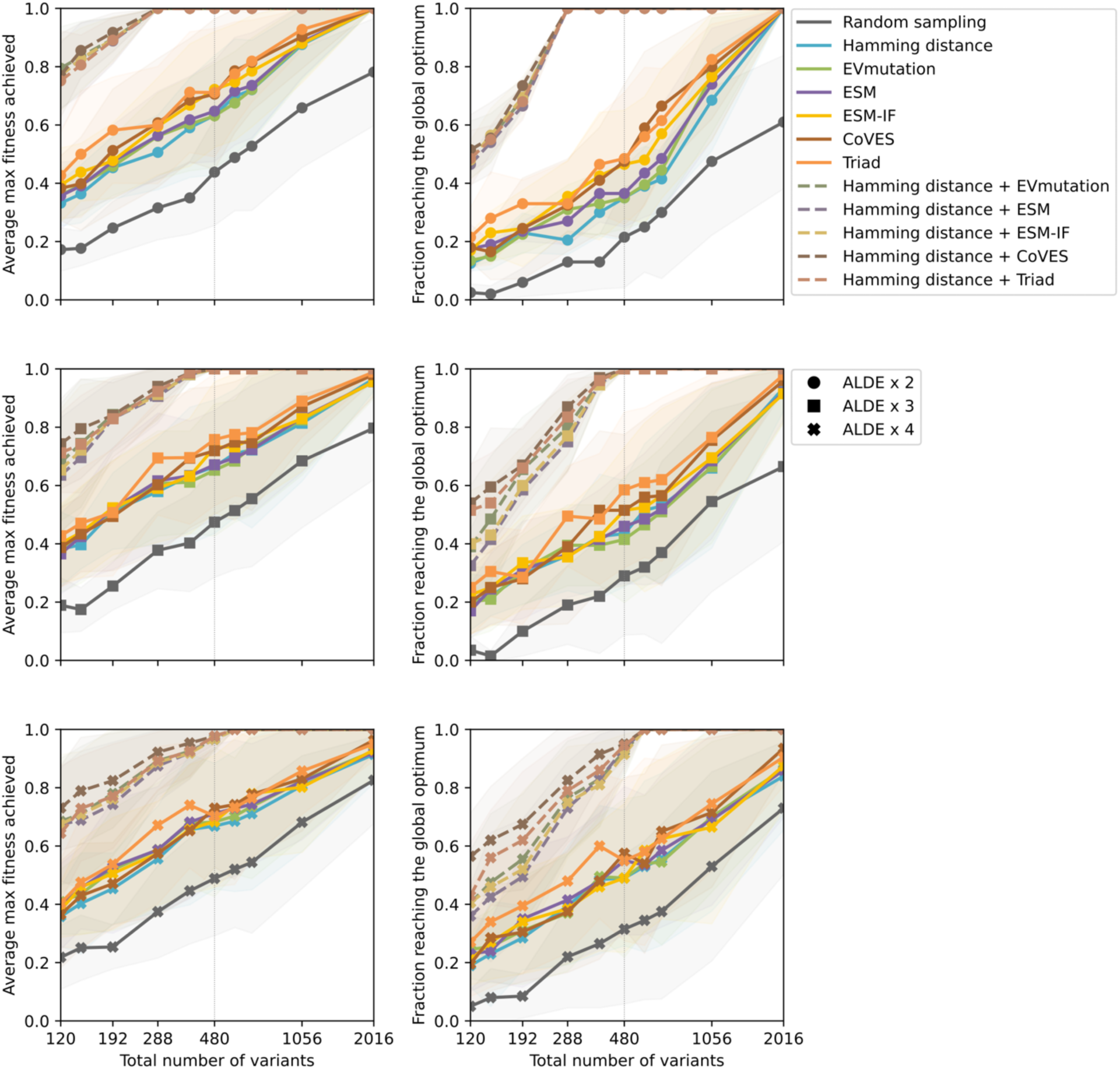
Multiple rounds of ftALDE averaged across four landscapes with fewer than 1% active variants, related to Figure 3e. Shading indicates standard deviation.

**Figure S10.**
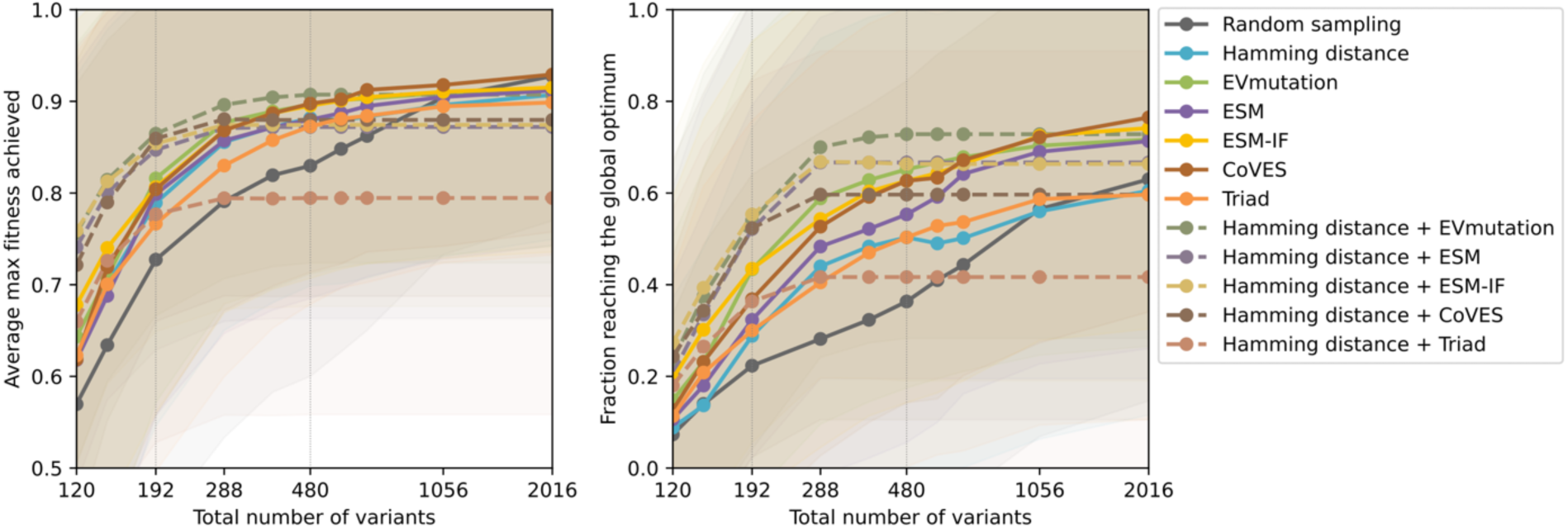
ftMLDE with Hamming distance-ensembled ZS predictors, averaged across 12 landscapes with more than 1% active variants, related to Figure 3e. Shading indicates standard deviation.

**Figure S11.**
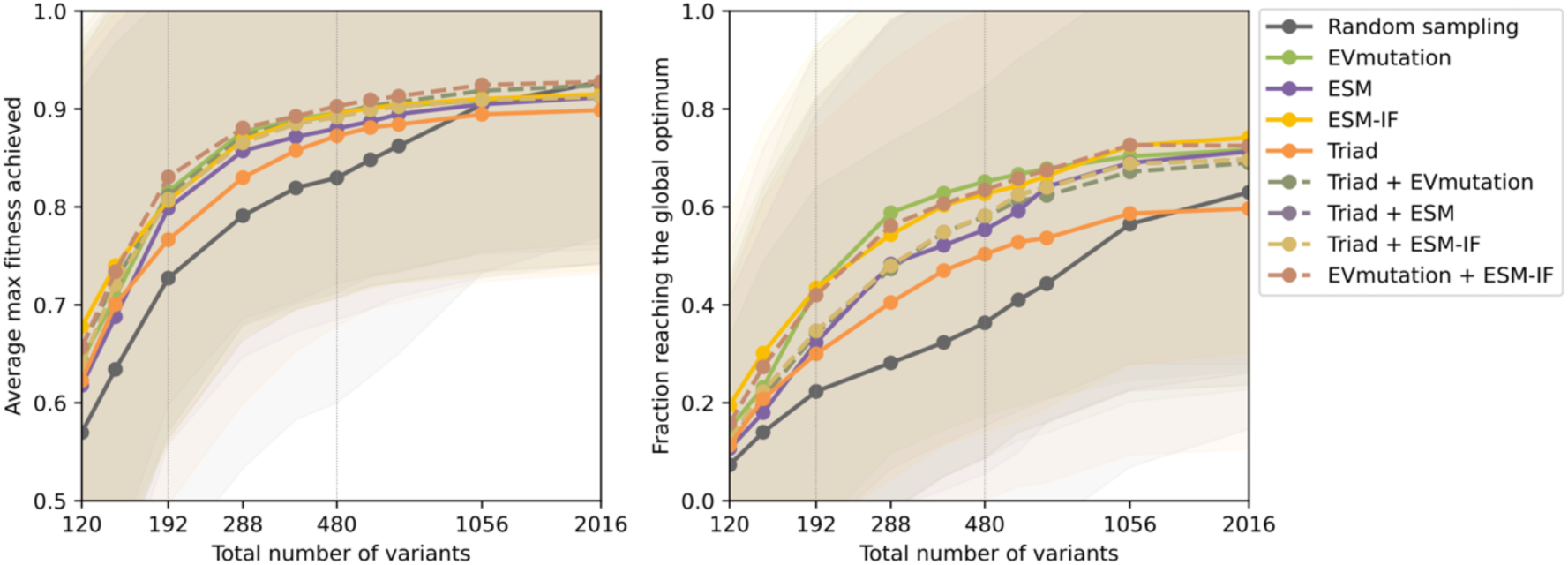
ftMLDE with Triad-ensembled ZS predictors or ESM-IF and EVmutation ensemble, averaged across 12 landscapes with more than 1% active variants, related to Figure 3e. Shading indicates standard deviation.

**Figure S12.**
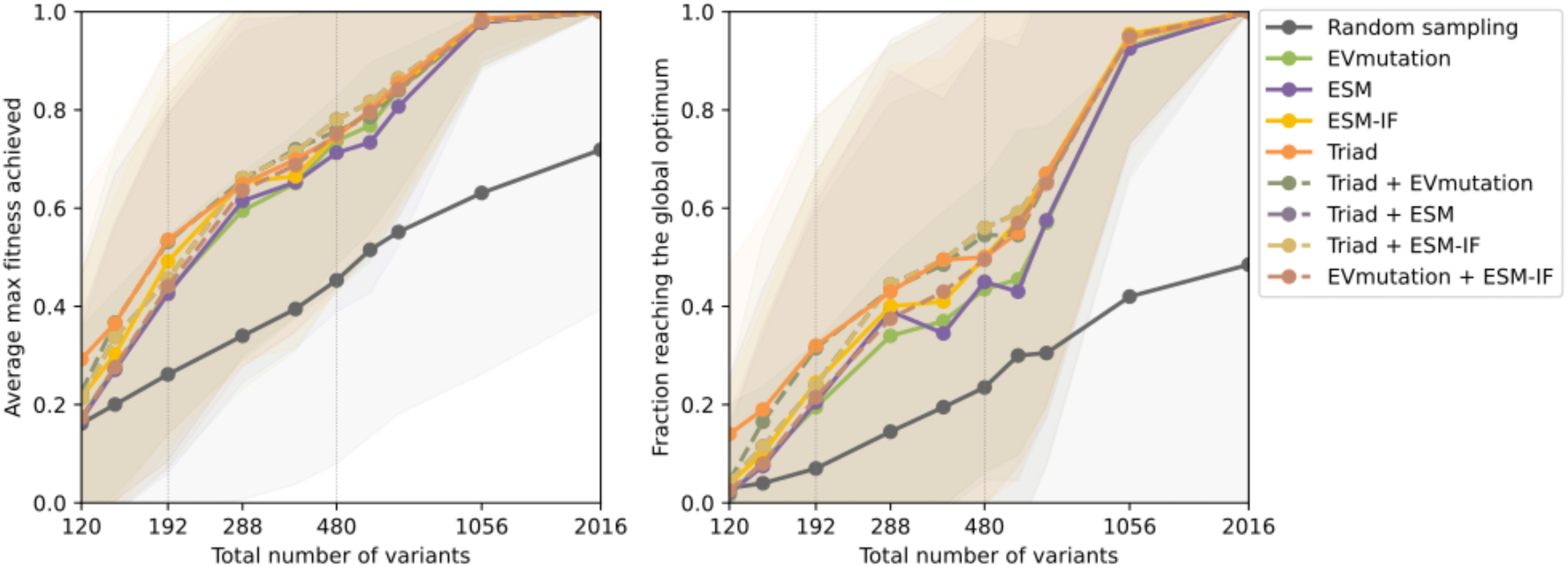
ftMLDE with Triad-ensembled ZS predictors or ESM-IF and EVmutation ensemble, averaged across four landscapes with fewer than 1% active variants, related to Figure 3e. Shading indicates standard deviation.

**Figure S13.**
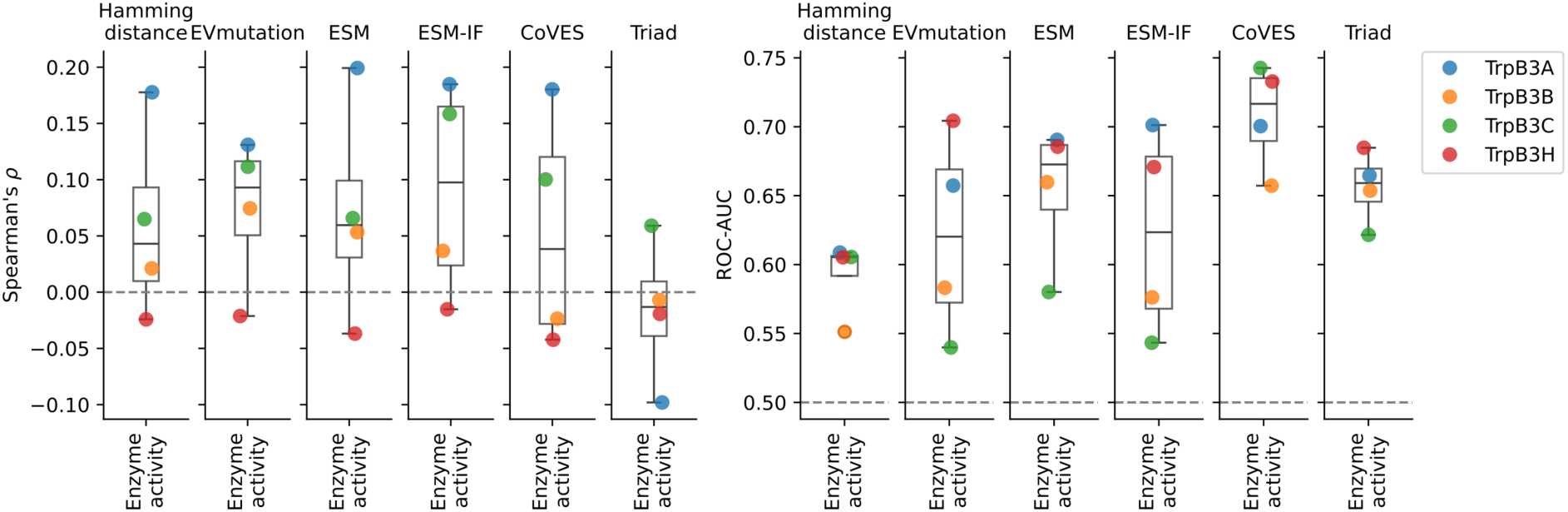
ZS predictor fitness value ranking (left) and active/inactive variant classification (right) for four landscapes with fewer than 1% active variants, related to Figure 4a.

**Figure S14.**
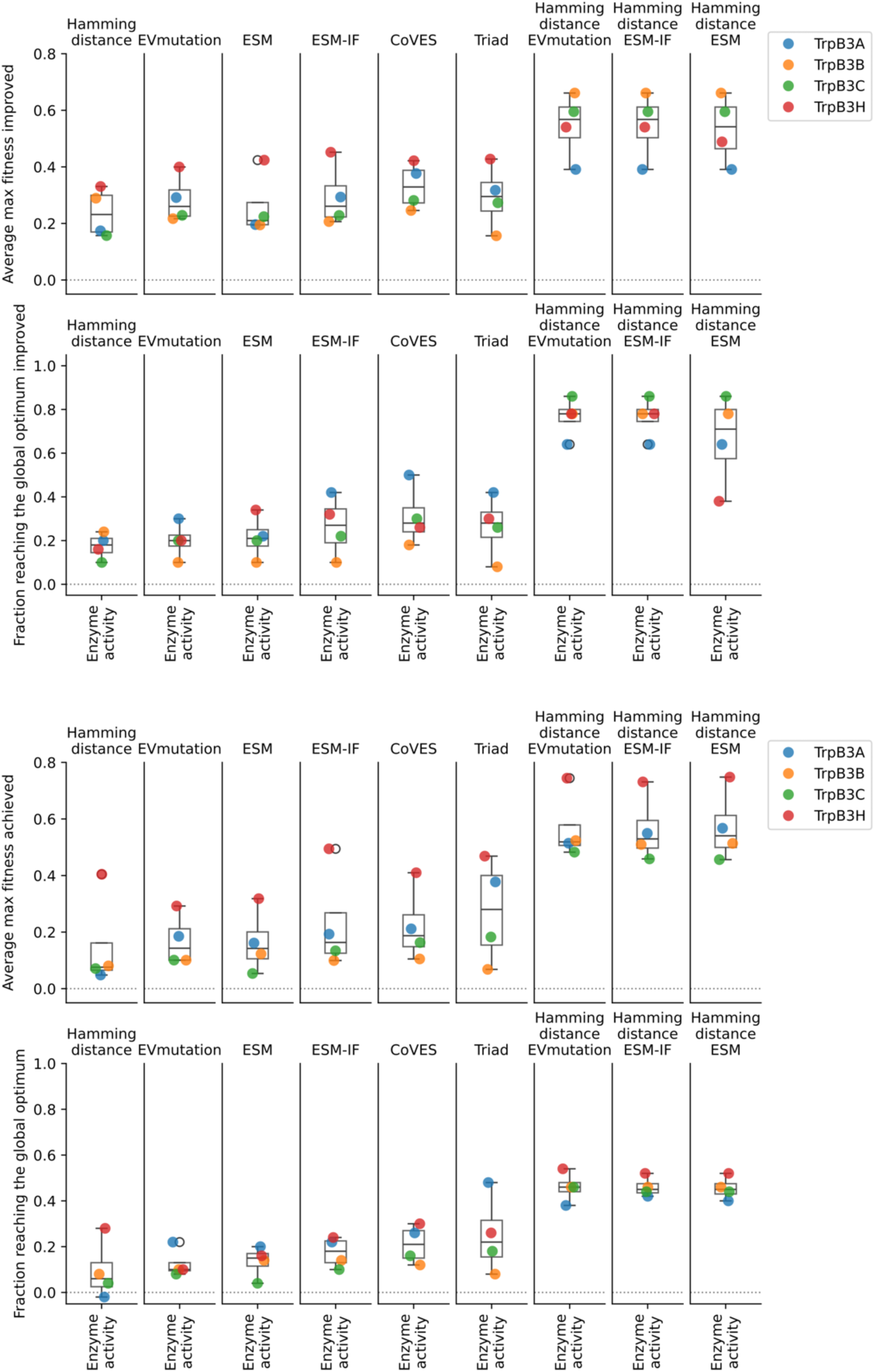
Effects of focused training for ftMLDE with a total sample size of 480 (384 training and 96 testing, top) and 192 (96 training and 96 testing, bottom) for four landscapes with fewer than 1% active variants, related to Figure 4b.

**Figure S15.**
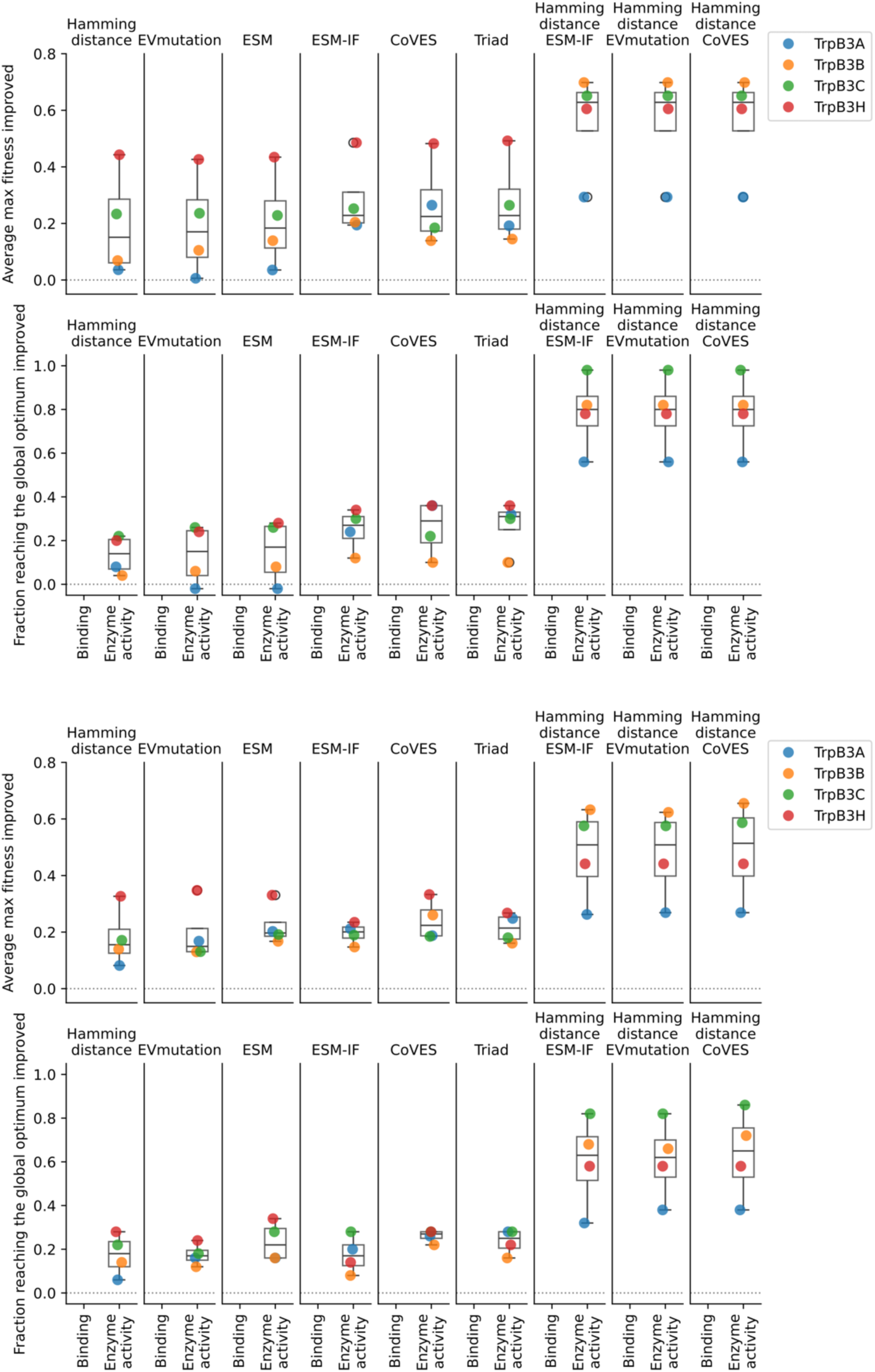
Effects of focused training for two (top) and four (bottom) rounds of ftALDE with a total sample size of 480 for four landscapes with fewer than 1% active variants, related to Figure 4b.

**Figure S16.**
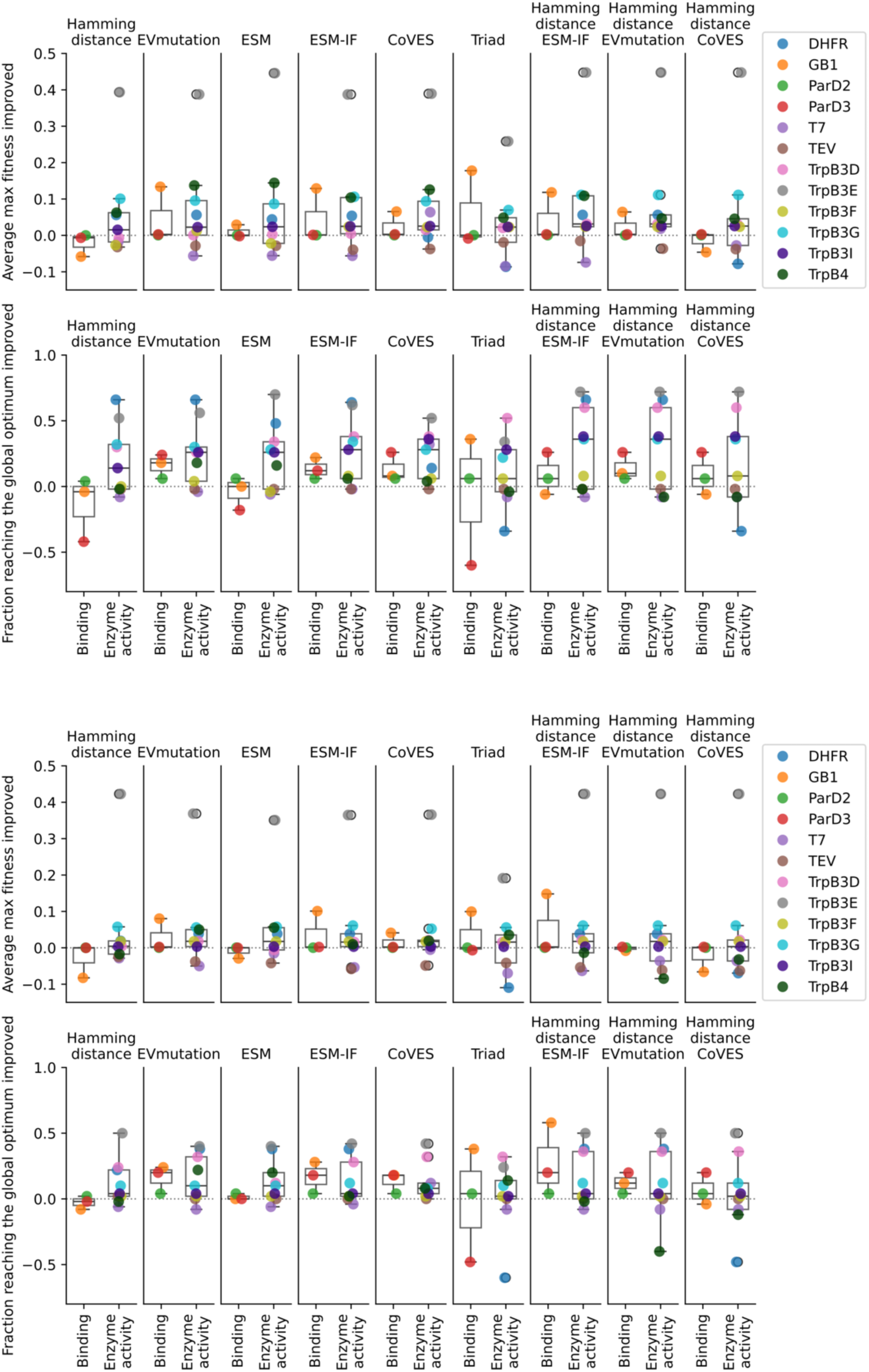
Effects of focused training for two (top) and four (bottom) rounds of ftALDE with a total sample size of 480 for 12 landscapes with at least 1% active variants, related to Figure 4b.

**Figure S17.**
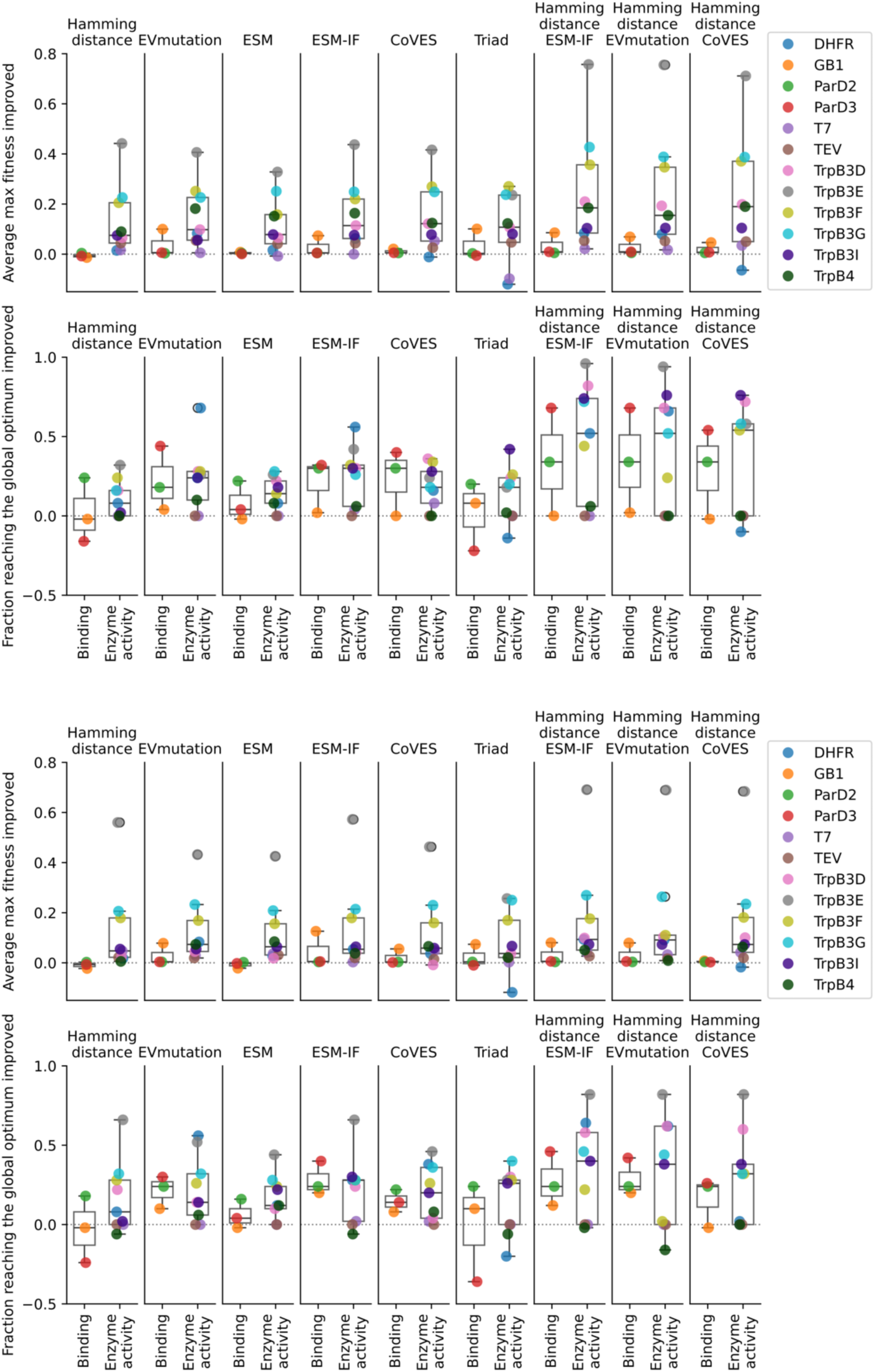
Effects of focused training for two (top) and four (bottom) rounds of ftALDE with a total sample size of 192 for 12 landscapes with at least 1% active variants, related to Figure 4b.

**Figure S18.**
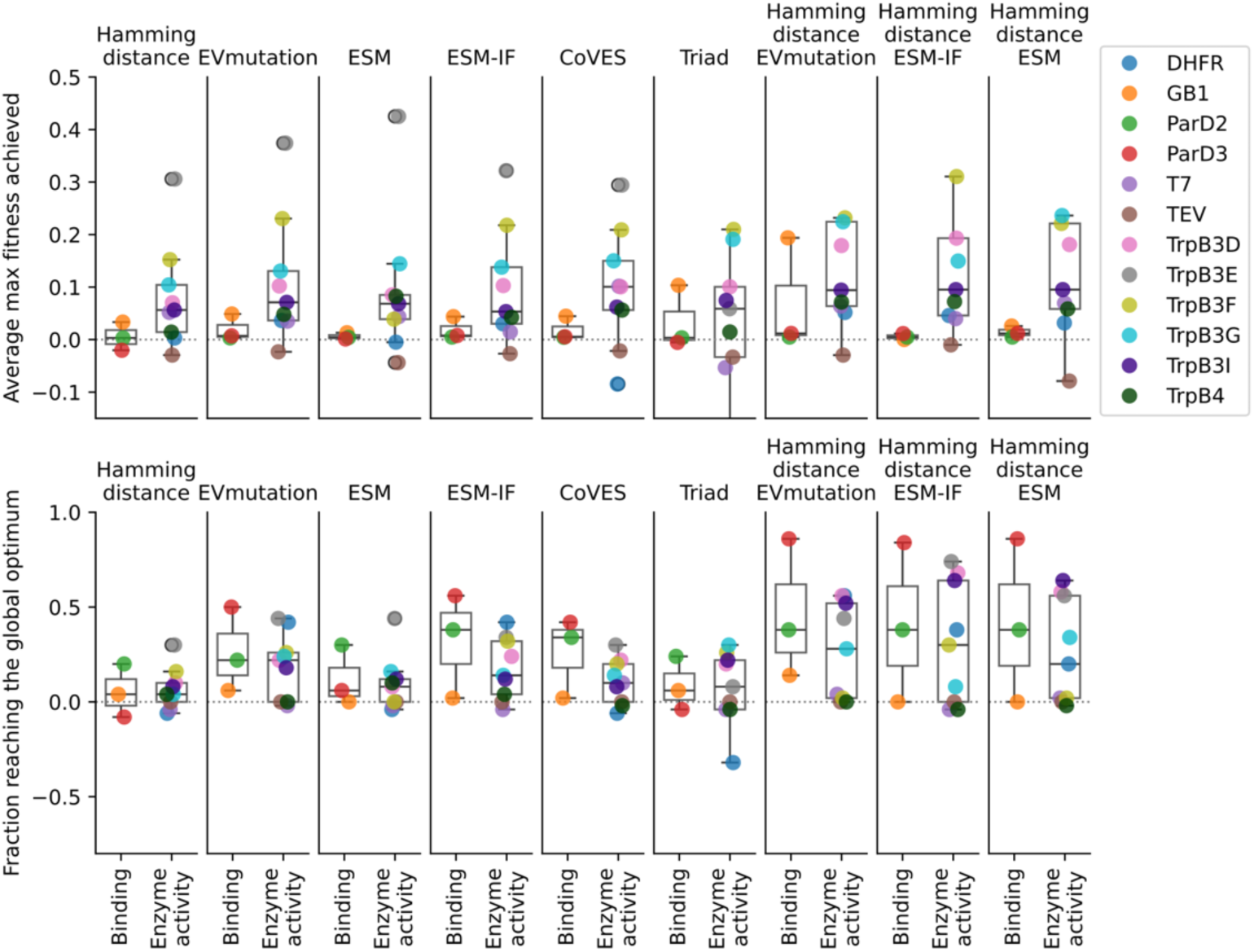
Effects of focused training for ftMLDE with a total sample size of 192 (96 training and 96 testing) for 12 landscapes with at least 1% active variants, related to Figure 4b.

**Figure S19.**
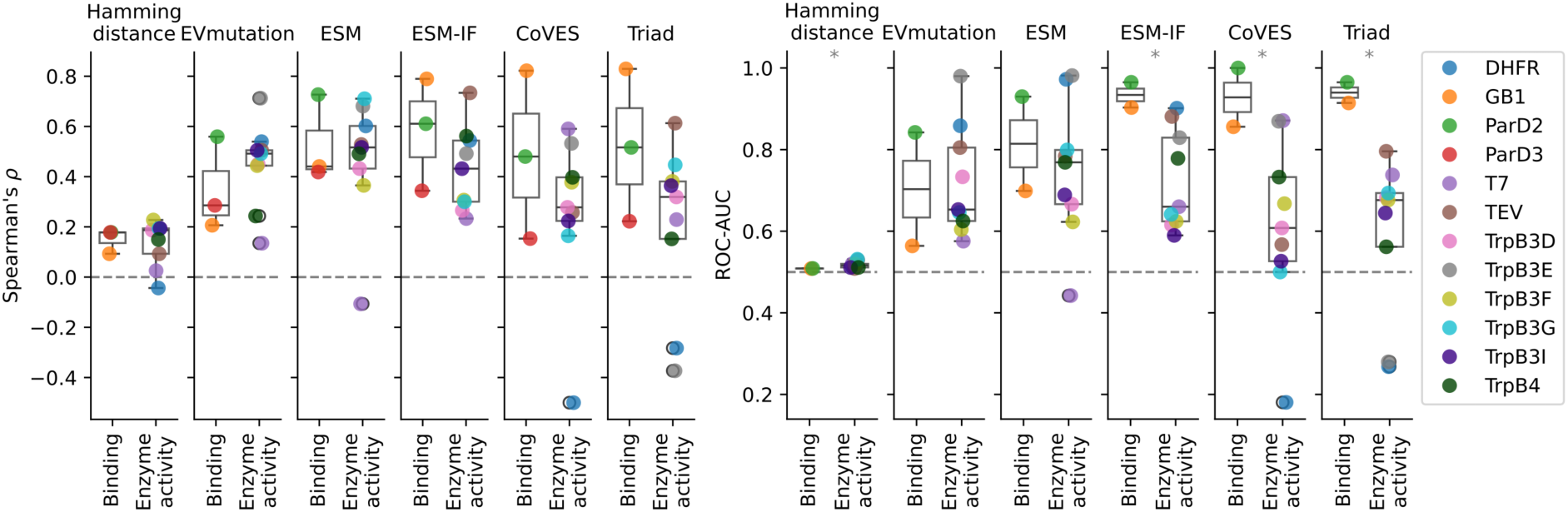
ZS predictor for single substitution fitness value ranking (left) and active/inactive variant classification (right) for 12 landscapes with at least 1% active variants, related to discussion and Figure 4a. Statistical significance (p-value <0.05) is indicated as *.

**Figure S20.**
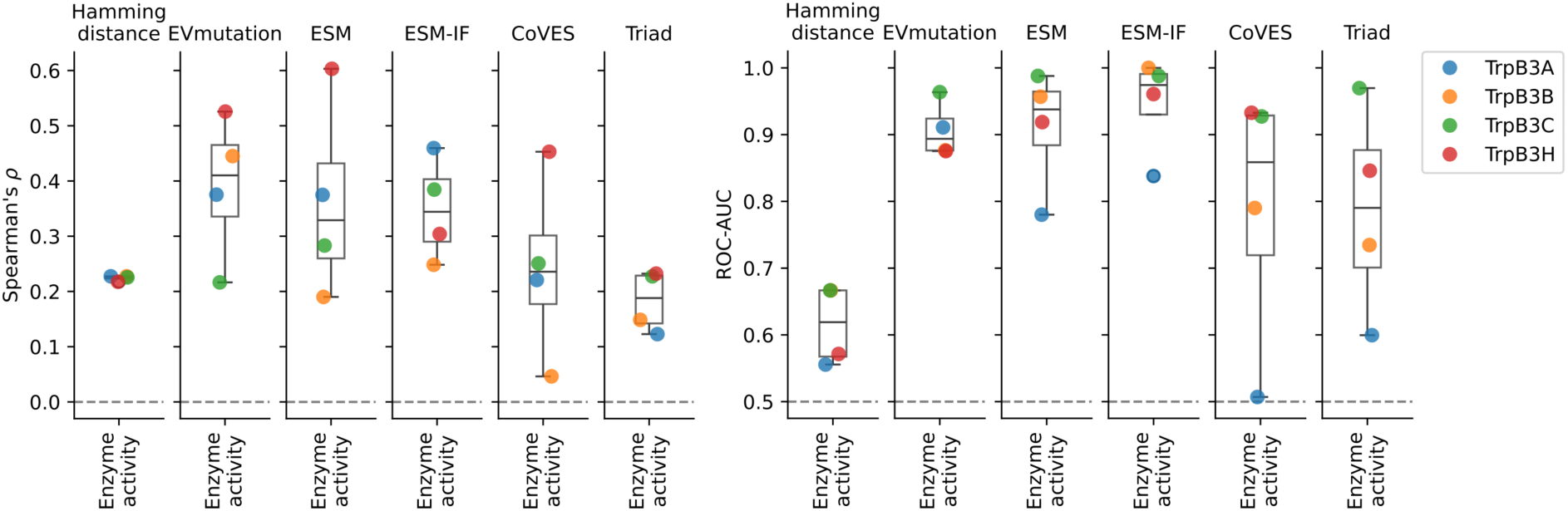
ZS predictor for single substitution fitness value ranking (left) and active/inactive variant classification (right) for landscapes with fewer than 1% active variants, related to discussion. Statistical significance (p-value <0.05) is indicated as *, related to discussion and Figure 4a.

**Figure S21.**
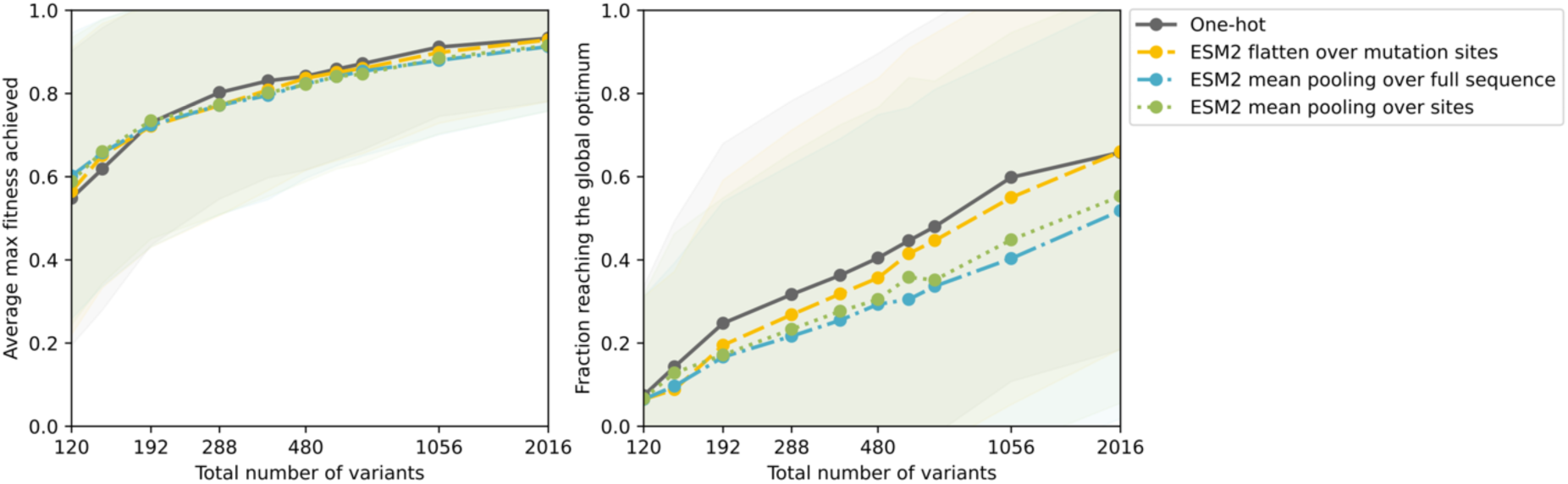
Encoding strategies for MLDE performance, averaged across 12 landscapes with at least 1% active variants. Comparison of learned embeddings from the protein language model ESM2 using different pooling methods vs. one-hot encoding flattened over the substitution sites, related to discussion. Shading indicates standard deviation.

**Figure S22.**
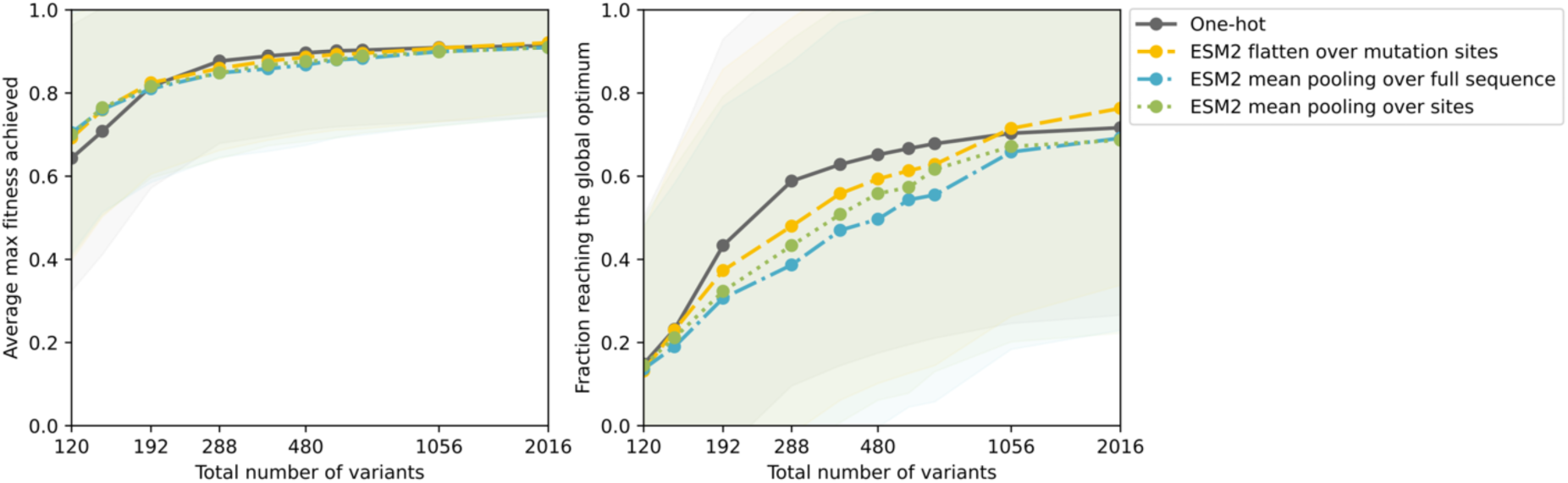
Encoding strategies for EVmutation-guided ftMLDE performance, averaged across landscapes with at least 1% active variants. Comparison of learned embeddings from the protein language model ESM2 using different pooling methods vs. one-hot encoding flattened over the substitution sites, related to discussion. Shading indicates standard deviation.

**Figure S23.**
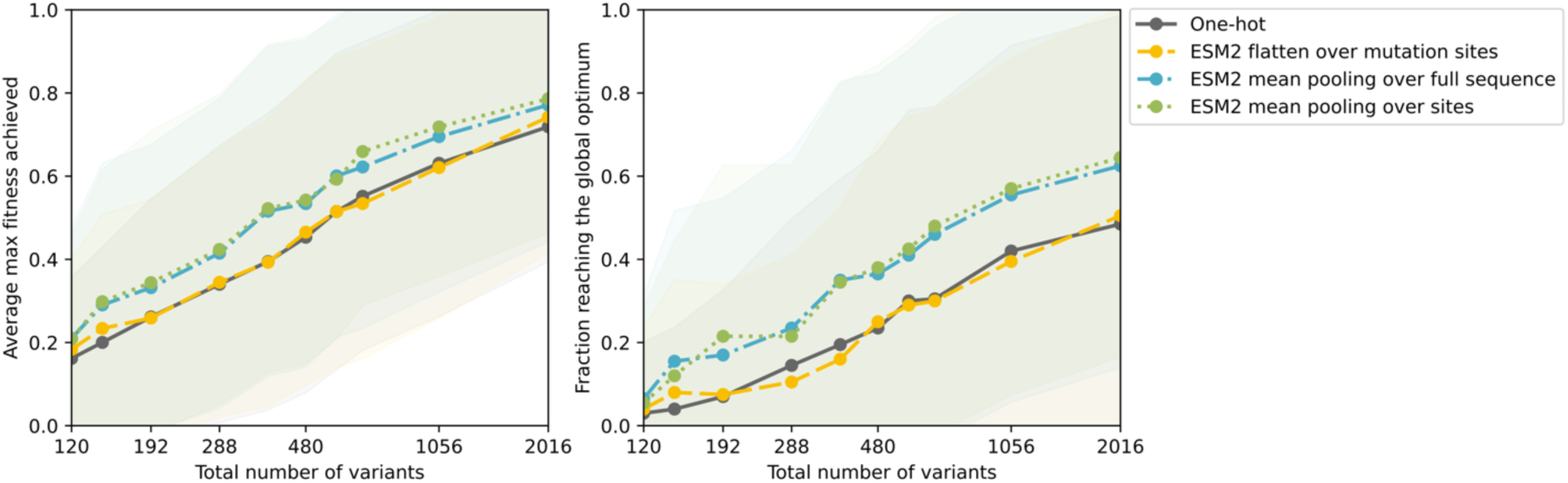
Encoding strategies for MLDE performance, averaged across four landscapes with fewer than 1% active variants. Comparison of learned embeddings from the protein language model ESM2 using different pooling methods vs. one-hot encoding flattened over the substitution sites, related to discussion. Shading indicates standard deviation.

**Figure S24.**
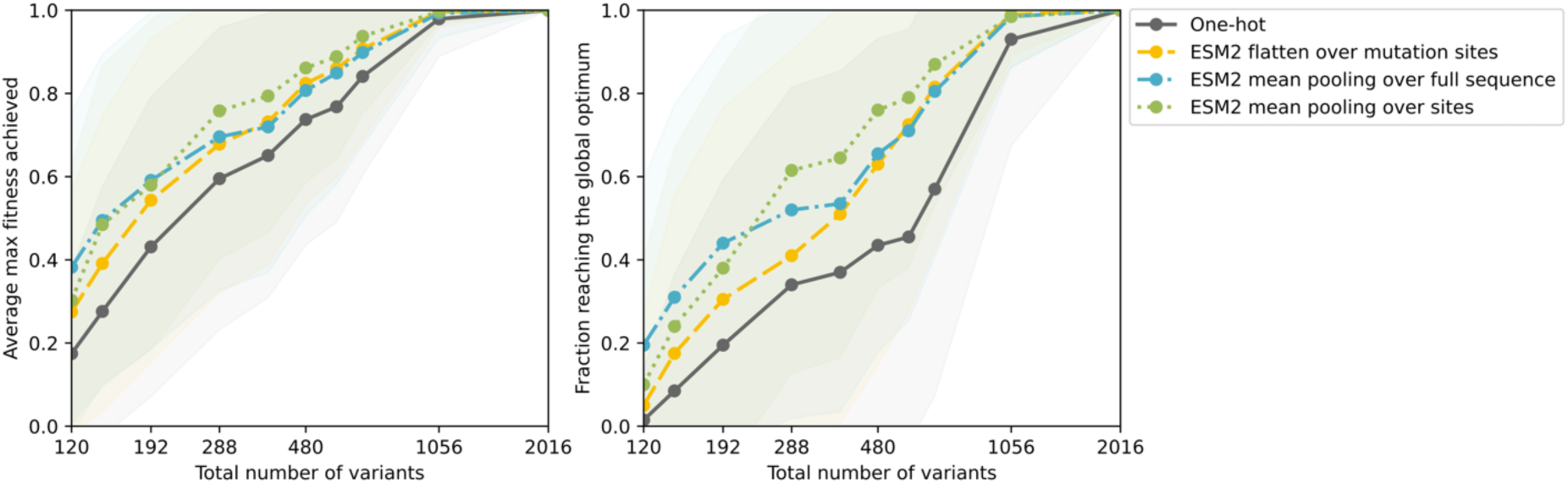
Encoding strategies for EVmutation-guided ftMLDE performance, averaged across four landscapes with fewer than 1% active variants. Comparison of learned embeddings from the protein language model ESM2 using different pooling methods vs. one-hot encoding flattened over the substitution sites, related to discussion. Shading indicates standard deviation.

**Figure S25.**
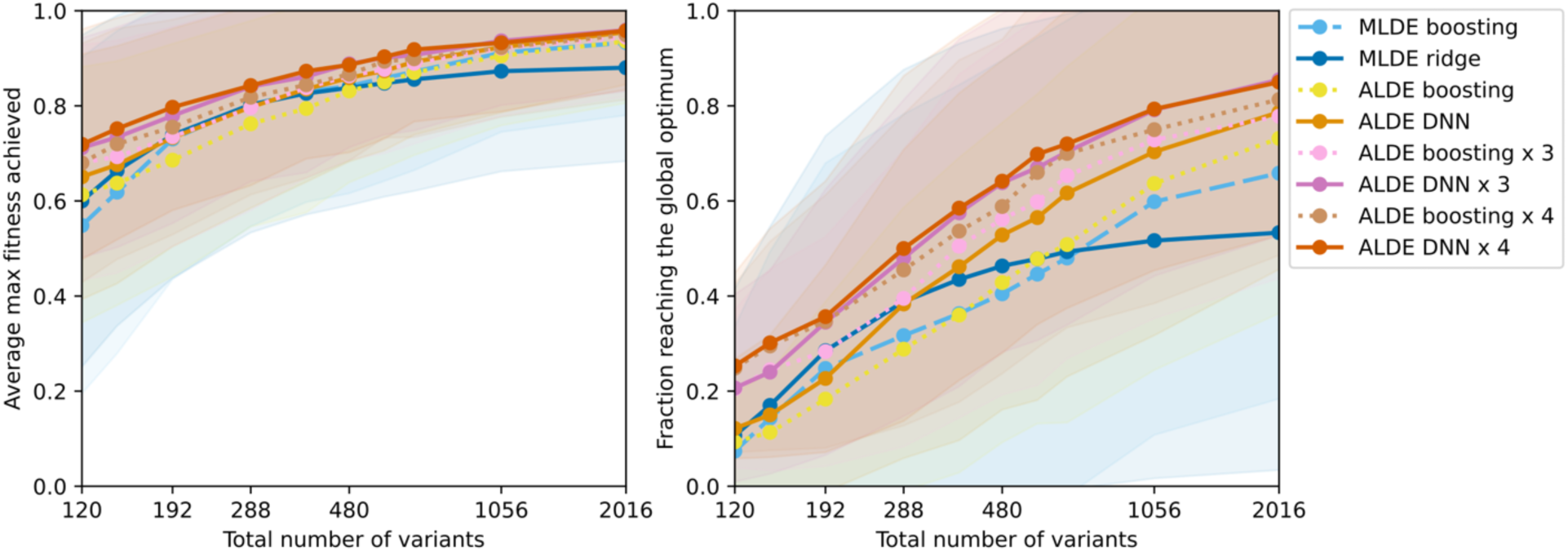
MLDE and ALDE with different model types, averaged across 12 landscapes with at least 1% active variants. MLDE with boosting or ridge regression. ALDE different rounds with boosting or deep neural network ensembles. No focused training included, related to discussion. Shading indicates standard deviation.

**Figure S26.**
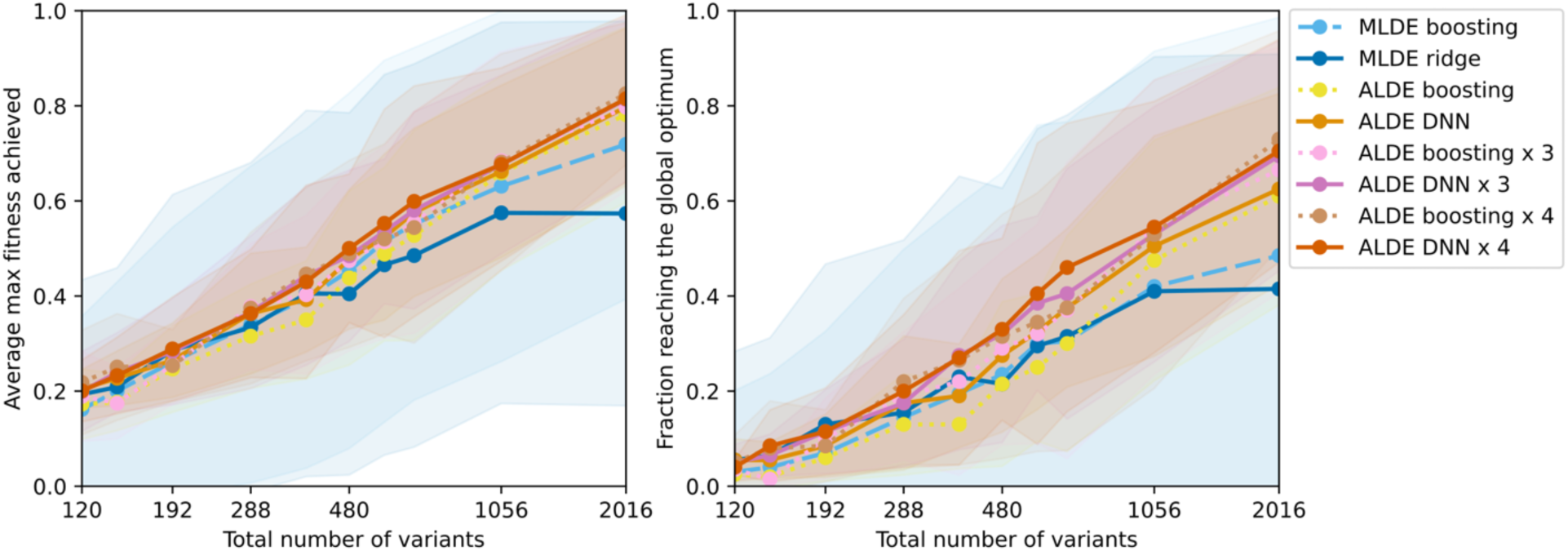
MLDE and ALDE with different model types, averaged across four landscapes with fewer than 1% active variants. MLDE with boosting or ridge regression. ALDE different rounds with boosting or deep neural network ensembles. No focused training included, related to discussion. Shading indicates standard deviation.

**Figure S27.**
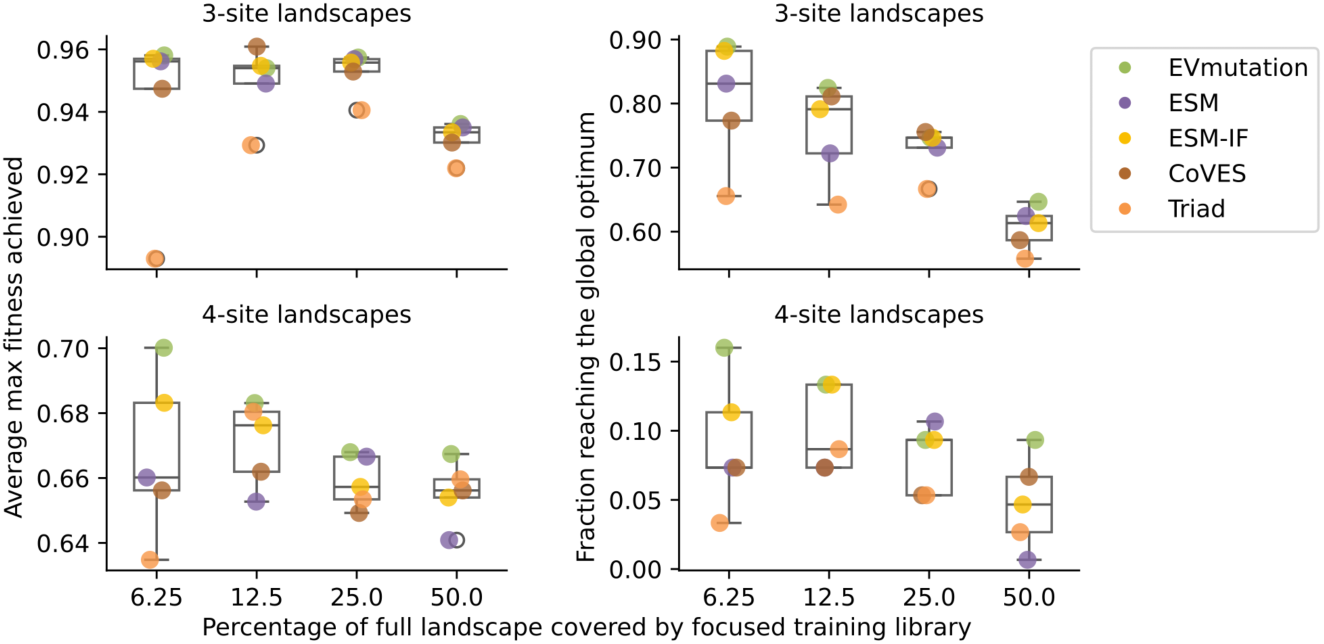
The impact of reducing the size of the focused training library relative to the full library on ftMLDE performance averaged across 12 landscapes with at least 1% active variants split into three-site landscapes (top row) and four-site landscapes (bottom row). Related to discussion.

